# CellCharter reveals spatial cell niches associated with tissue remodeling and cell plasticity

**DOI:** 10.1101/2023.01.10.523386

**Authors:** Marco Varrone, Daniele Tavernari, Albert Santamaria-Martínez, Logan A. Walsh, Giovanni Ciriello

**Affiliations:** Department of Computational Biology, University of Lausanne, Lausanne, Switzerland; Swiss Cancer Center Léman, Lausanne, Switzerland; Swiss Institute of Bioinformatics, Lausanne, Switzerland; Swiss Institute for Experimental Cancer Research, École Polytechnique Fédérale de Lausanne (EPFL), Lausanne, Switzerland; Rosalind and Morris Goodman Cancer Institute, McGill University, Montreal, Quebec, Canada; Department of Human Genetics, McGill University, Montreal, Quebec, Canada

## Abstract

Tissues are organized in cellular niches, the composition and interactions of which can be investigated using spatial omics technologies. However, systematic analyses of tissue composition are challenged by the scale and diversity of the data. Here we present CellCharter, an algorithmic framework to identify, characterize, and compare cellular niches in spatially resolved datasets. CellCharter outperformed existing approaches and effectively identified cellular niches across datasets generated using different technologies, and comprising hundreds of samples and millions of cells. In multiple human lung cancer cohorts, CellCharter uncovered a cellular niche composed of tumor-associated neutrophils and cancer cells expressing markers of hypoxia and cell migration. This cancer cell state was spatially segregated from more proliferative tumor cell clusters and was associated with tumor-associated neutrophil infiltration and poor prognosis in independent patient cohorts. Overall, CellCharter enables systematic analyses across data types and technologies to decode the link between spatial tissue architectures and cell plasticity.

## MAIN

High-throughput molecular profiling allows one to decipher heterogeneity and biological activity among species, organs, and cells. Technological advancements in the past few decades have enabled the characterization of molecular features unbiasedly, often at single-cell resolution^1^. Spatial omics approaches represent the latest revolution in this progress^2^. These technologies quantify molecular features in single cells while retaining information about the physical location of each cell within the tissue^3,4^. Spatial molecular profiles have been shown to recapitulate tissue anatomy and reveal cellular niches characterized by specific admixtures of different cell types^5–7^. In cancer, these technologies have been used to investigate the association between cellular patterns and disease aggressiveness^8–11^, or the oncogenic potential of mutated cells based on their location^12,13^. However, these technologies are still limited in terms of resolution and/or scalability. Currently, spatial proteomics and transcriptomics approaches mostly differ in terms of *resolution* and *coverage*. Spatial proteomics largely relies on multiple cycles of multiplexed immunofluorescence or on imaging mass cytometry/spectrometry^3,14^. As such, they provide single-cell resolution data but low coverage, quantifying only tens or a few hundred proteins. Spatial transcriptomics can be divided into image-based and sequencing-based approaches^15,16^. The former assays up to 1,000 genes but achieves single-cell or even subcellular resolution. The latter can cover the entire transcriptome but within fixed-size spots that comprise between 10 and 100 cells. High-resolution implementations of sequencing-based spatial transcriptomics have been proposed but they have not yet reached standardization and commercialization^17–21^. Recently, the development of multiome spatial technologies allow to simultaneously assay multiple molecular features within the same cell or spot^22^.

With the development of new technologies, new computational approaches have emerged to analyze spatial molecular profiles. Among these, *spatial clustering* assigns cells to clusters based on both their intrinsic features, such as protein or messenger RNA abundance, and the features of neighboring cells in the tissue. Hence, whereas classical clustering determines populations of molecularly similar cells, spatial clustering determines cellular *niches* characterized by the admixture of specific cell populations. A simple approach to determine cellular niches is clustering cells based on the proportion of cell types within their neighborhood^5,23,24^. Systematic approaches for spatial clustering include Hidden Markov Random Fields (HMRFs^25^), which extend Hidden Markov Models to an unbounded number of neighbors, or Graph Neural Networks (GNNs^26^), a machine learning technique that generates representations of a node (e.g., a cell) from the convolution of features of its neighbors in a network^27^. A Bayesian version of HMRF is used in BayesSpace^28^, whereas DR-SC^29^ combines dimensionality reduction of the feature space and HMRF. SpaGCN^30^, SEDR^31^ and STAGATE^32^ use a GNN to determine network representations of spatial datasets. SPACE-GM uses a GNN to perform supervised learning on patient clinical outcome^33^. Without resorting to a GNN, UTAG annotates a cell by its features and the features of its direct neighbors^34^. Recent alternative approaches for spatial clustering are SpatialPCA^35^, which uses a kernel matrix in probabilistic Principal Component Analysis (PCA) to model spatial correlations, and SOTIP^36^, which clusters cells based on a network of cell neighborhoods.

Outstanding challenges in the field concern scalability and portability. Recent large-scale applications of image mass cytometry (IMC) assays^23,24,37,38^ or the development of technologies profiling large tissue slides at single-cell resolution^39–41^ require tools capable of scaling with large numbers of both samples and cells. However, most methods have not been designed to simultaneously cluster, characterize, and compare large numbers of samples or cells. Beyond scalability, clustering multiple samples requires correcting for potential batch effects. Currently, only BayesSpace and UTAG include a batch effect correction procedure. Moreover, with rapidly evolving technologies, tools need to be usable with data generated by different techniques. None of the current approaches satisfies this criterion, either because they require specific cell layouts or because they cannot scale with high-throughput single-cell assays. Even more, with the advent of spatial multiome platforms, approaches able to integrate different data modalities are highly desirable. Finally, beyond the identification of spatial clusters, discoveries from spatially resolved datasets will come from the characterization and comparison of such clusters across tissue types and conditions. Here, we introduce CellCharter (https://github.com/CSOgroup/cellcharter), a new algorithmic framework to address these challenges.

## RESULTS

### Inference, characterization and comparison of cell niches

We designed CellCharter to: 1) analyze large cohorts of spatially profiled samples; 2) be agnostic of the underlying technology used to generate spatial molecular profiles; and 3) implement not only a spatial clustering algorithm, but also a suite of approaches for cluster characterization and comparison. CellCharter takes as input a spatial omics dataset represented by a matrix of features; for example, mRNA or protein abundance in a cell or spot, along with the spatial coordinates of each cell/spot (**Fig. 1a** - left). Dimensionality reduction and batch effect correction are then performed using variational autoencoders^42–44^ (VAEs), which are neural networks mapping the input features into embeddings in a latent space. Because different omics assays often have distinct data distributions, we select an appropriate VAE for each data type (**Methods** and **Supplementary Methods**). Next, CellCharter constructs a network of cells/spots based on their spatial proximity and, for each cell/spot *A*, we define the *l*-neighborhood of *A* as the set of cells/spots that are at most *l* steps away from *A* in the network, where *l* is a user-defined parameter (in **Fig. 1a** - center, *l* = 3). To aggregate the features of each cell/spot with those of its *l*-neighborhood, we concatenate the feature vector of *A* and *l* vectors, each containing the feature averages of cells/spots at *i* steps from *A*, for each *i* ∈ [1, *l*]. Lastly, cells/spots are clustered based on this concatenated vector of aggregated features using a Gaussian Mixture Model (GMM). To determine the number of clusters, GMM is run multiple times (n = 10 in this study) for each possible number of clusters within a user-defined range. A solution with *n* clusters is considered “stable” when cluster assignments are reproducible across multiple GMM runs for *n*, *n - 1*, and *n + 1*, based on the Fowlkes-Mallows Index^45^ (FMI; **Fig. 1a** - right). By incorporating a batch effect correction step and using a highly scalable approach to encode spatial information, CellCharter is particularly suited to simultaneously determine spatial clusters among multiple samples.

**Figure 1:**
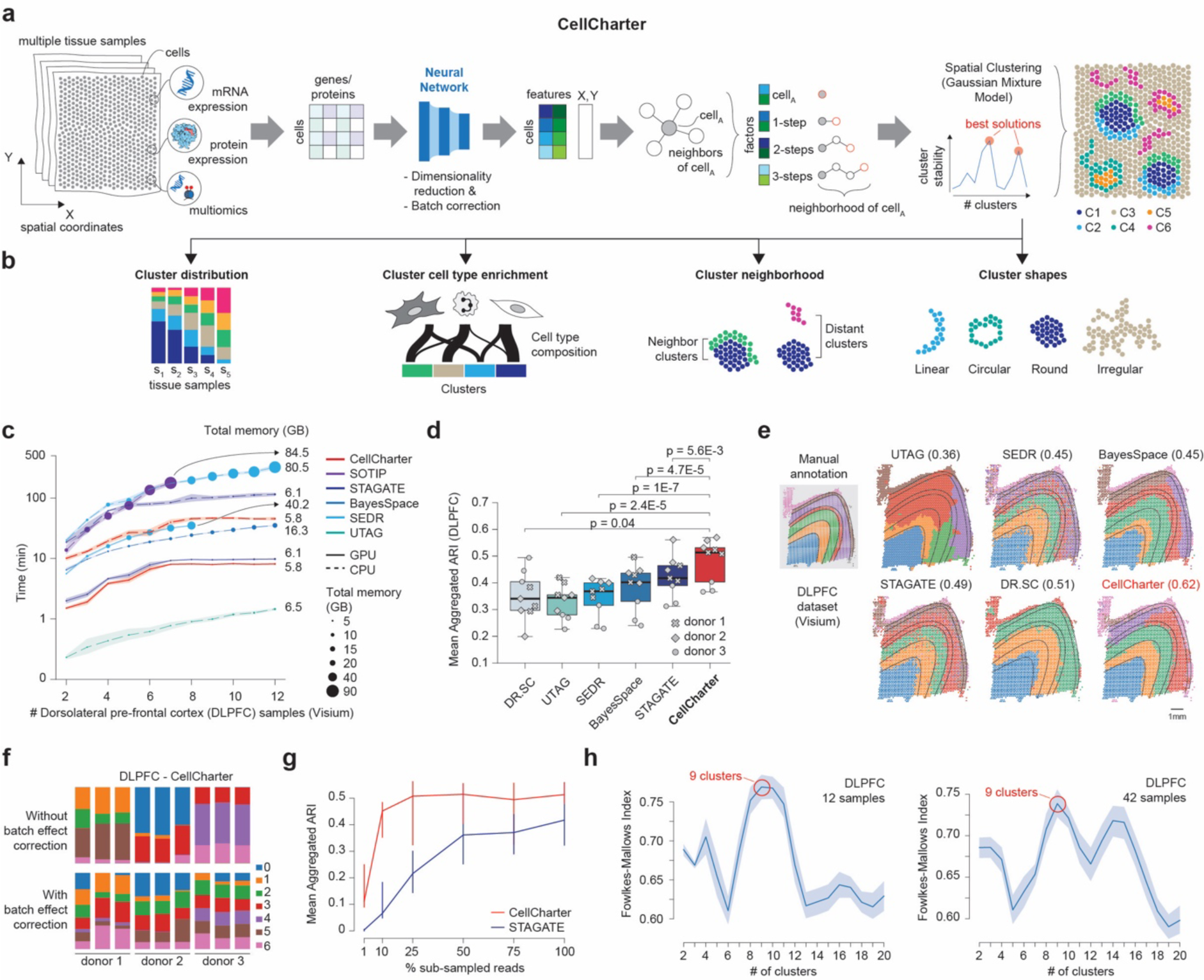
CellCharter identifies, characterizes, and compares spatial clusters. **a)** Workflow of CellCharter. From left to right: spatial molecular profiling generates a gene, protein and/or multiomics expression matrix comprising the coordinates (x,y) of each cell. Dimensionality reduction and batch correction are performed on this matrix and, for each cell (cell_A_), its features are concatenated with the features of its neighbors. Lastly, Gaussian Mixture Model clustering is performed and best cluster solutions are chosen based on cluster stability. **b)** Downstream analyses implemented in CellCharter for the characterization and comparison of spatial clusters. **c)** Runtime and memory requirement for the evaluated clustering methods at increasing DLPFC dataset size. Runtime is presented as mean values with a 95% confidence interval. **d)** Mean Adjusted Rand Index (ARI) for each DLPFC sample (over 10 repetitions, y axis) obtained by the listed methods (x axis) upon performing joint spatial clustering of all test samples (n = 9). Boxes show the quartiles of the dataset, whereas whiskers extend to points that lie within 1.5 inter-quartile ranges (IQRs) of the lower and upper quartile. p-values were computed by two-sided t-test. **e)** Manually annotated and predicted cluster labels by all the evaluated methods for one tissue slide from the Visium DLPFC dataset upon performing joint spatial clustering of all test samples (n = 9). The best ARI value of 10 repetitions is reported. **f)** CellCharter spatial cluster distribution across 9 DLPFC samples without batch effect correction (upper) and with batch effect correction (lower). **g)** Mean aggregated ARI values obtained by CellCharter (red) and STAGATE (blue) at different sequencing depths (mean ARI over 10 runs per condition) on all test samples (n = 9). Error bars correspond to the lower and upper 95^th^ percentiles. **h)** CellCharter cluster stability (y axis) for range of number of clusters (x axis) for the pilot DLPFC cohort (n = 12 samples) and the extended DLPFC cohort (n = 42 samples). Most stable cluster solution is highlighted. Data are presented as mean values with a 95% confidence interval.

In addition, CellCharter implements downstream analyses to: (1) determine cluster proportions for each sample; (2) compute cell type enrichment for each cluster (when cell type annotations are available); (3) estimate significant spatial proximity among clusters (cluster neighborhood enrichment or cluster NE) and how it varies among conditions (*differential* cluster NE); and (4) characterize and compare cluster *shapes* (**Fig. 1b**). CellCharter introduces an analytical approach to compute *symmetric* and *asymmetric* cluster NE, which is more efficient than currently available permutation-based methods^46^. Unlike in the symmetric case, asymmetric cluster NE allows discriminating when the neighborhood of one cluster is enriched for another cluster but not vice versa (**Supplementary Figure Fig. 1a**). In addition, CellCharter characterizes the shape of each *cluster component*, that is, the set of cells belonging to the same cluster and within a connected component of the cell network that CellCharter uses to encode spatial proximity. For each cluster component, CellCharter computes its *boundary*, from which it derives the area and perimeter of the component, and its *bounding box*, i.e., the smallest rectangle encasing the cluster component, from which it derives the minor and major axes (**Methods**). Based on these measures, we compute four scores: **curl**, which expresses how curved or twisted a shape is; **elongation**, which is the ratio of the major and minor axes; **linearity**, which assesses how well a shape can be approximated by a linear path; and **purity**, which quantifies the fraction of cells within a cluster boundary that belongs to that cluster (**Supplementary Figure Fig. 1b**). Representative cluster components from a spatial proteomics dataset generated by CODEX^5^ demonstrate how combinations of these scores allow to describe shapes as linear, round, circular and irregular (**Supplementary Figure Fig. 1c**).

To demonstrate the effectiveness and efficiency of CellCharter, we first assessed its spatial clustering performance against state-of-the-art approaches. For this purpose, we used an annotated spatial transcriptomics dataset (10x Genomics Visium) comprising 12 samples (4 samples each from 3 donors) of human dorsolateral prefrontal cortex (DLPFC)^47^. Within each sample, spots have been manually assigned to one of six cortical layers (L1-L6) or to white matter (WM) (**Supplementary Figure Fig. 1d**), which are shared among samples and were used as ground truth. Based on previous benchmarking analyses^28,29,31,32,31^, we compared CellCharter with 7 different approaches, testing their running time, memory usage and cluster quality, assessed using the Adjusted Rand Index (ARI). To avoid performing hyperparameter tuning on the samples used for testing (**Supplementary Figure Fig. 1e**), we introduced a separate tuning step using 3 samples (one for each donor) and then clustered the spots of the remaining 9 samples (**Supplementary Figure Fig. 1f** and **Supplementary Note**). Clustering was performed both on each individual sample and, when possible, jointly on all of them (**Methods**). SOTIP and SEDR (GPU version) could not jointly cluster all Visium samples because of their memory requirements. CellCharter exhibited the lowest memory usage, both in its GPU and CPU versions, and its computational efficiency was second only to BayesSpace, which was slightly faster although requiring more memory, and UTAG, which was the fastest approach (**Fig. 1c**), at the expense of low clustering quality (**Fig. 1d-e, Supplementary Figure Fig. 1g-h**). DR-SC was excluded from the runtime and memory comparison because it does not allow inclusion of a batch correction step, hence its execution time cannot be compared with the others. Importantly, without batch correction, clusters were associated with donors rather than tissue anatomy (**Fig. 1f**). Although STAGATE obtained the best cluster quality when clustering individual samples (**Supplementary Figure Fig. 1g**), CellCharter outperformed existing tools when jointly clustering all samples, both in terms of average ARI (**Fig. 1d, Supplementary Figure Fig. 1h**) and best ARI (**Fig. 1e**) over multiple runs. CellCharter was significantly faster than STAGATE, largely because of its efficient clustering procedure (**Supplementary Figure Fig. 2a**), and was robust to sequencing depth, maintaining similar high-quality performances upon retaining as little as 10% of the total amount of reads (**Fig. 1g**). Lastly, whereas all comparisons were made requiring each tool to detect 7 clusters, we tested our approach for the automatic selection of the number of clusters. CellCharter indicated n = 9 as the optimal number of clusters, close to the number of manually annotated regions, and the same number was found when clustering an extended version of this cohort comprising a total of 42 samples (**Fig. 1h**). By contrast, frequently used approaches such as Akaike Information Criterion (AIC)^48^, Bayesian Information Criterion (BIC)^49^, and Negative Log-Likelihood (NLL) either did not show a clear elbow or suggested a higher number of clusters, which increased upon increasing the number of samples to be clustered (**Supplementary Figure Fig. 2b**).

To further evaluate the performances of the best methods from previous comparisons, CellCharter and STAGATE, we used a single-cell resolution spatial proteomics dataset (CODEX platform) comprising 3 samples of normal mouse spleen and 6 samples of mouse spleen derived from an animal model of systemic lupus erythematosus^5^. Manual annotations from the original work classified regions in these samples as B follicles, marginal zones, periarteriolar lymphoid sheath (PALS), or red pulp (**Fig. 2a**). Spatial clustering was jointly performed on all samples. Visual comparisons of spatial clusters and manually annotated regions showed that CellCharter clusters best approximated the true tissue components (**Fig. 2a**), generating clusters with higher purity (**Supplementary Figure Fig. 2c**) and better mimicking the spleen anatomy than STAGATE. On this large dataset (707,466 cells in total), CellCharter was four times faster than STAGATE, largely because of its more scalable approaches for spatial features encoding and clustering (**Fig. 2b**). Finally, we tested whether embeddings generated from different data types could be integrated to cluster spatial multiome data^22^. We used CellCharter to perform spatial clustering either on single data types (RNA-seq or ATAC-seq) or on both, by using dedicated VAEs for embedding generation and joining these embeddings to define cell features. On this dataset, CellCharter identified 10 spatial clusters that better recapitulated the tissue anatomy when jointly using both data modalities than when using either of them alone^50^ (**Fig. 2c** and **Supplementary Figure Fig. 2d**).

**Figure 2:**
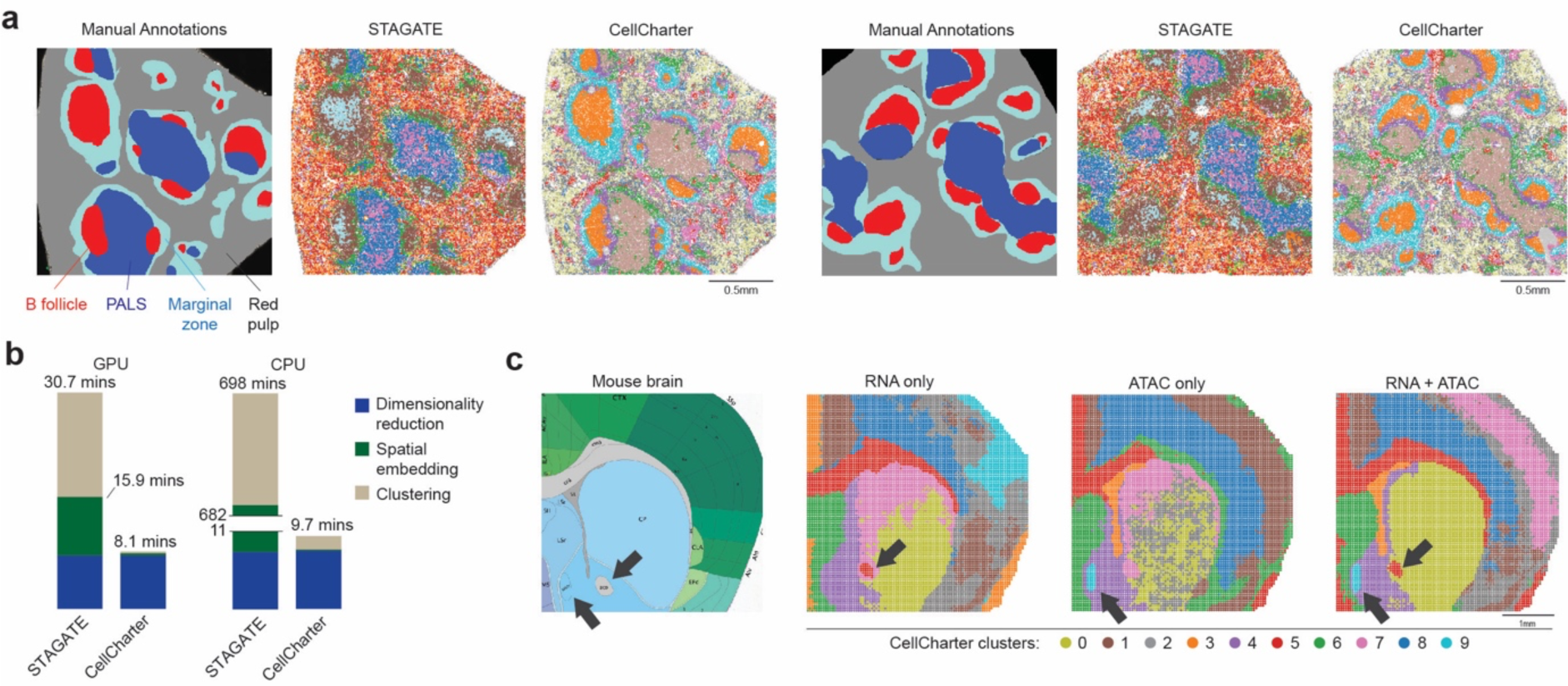
CellCharter is accurate, scalable and flexible. **a)** Manual annotations and spatial clusters (color coded) determined by STAGATE and CellCharter in two tissue samples of mouse spleen analyzed by CODEX. **b)** Running times of STAGATE and CellCharter to perform spatial clustering of 9 mouse spleen samples analyzed by CODEX. For both methods, the running time of the three main steps are shown separately. **c)** From left to right: manual annotations of the mouse brain (source: [Allen Reference Atlas – Mouse brain](http://atlas.brain-map.org/)), and CellCharter spatial clusters obtained based on spatially resolved RNA-seq (left), ATAC-seq (center) or multiome RNA + ATAC-seq (right) data. Arrows point to examples of clusters which are concurrently retrieved only in the multiome settings.

### Cellular niches in healthy and systemic lupus mice

The mouse spleen spatial proteomics dataset analyzed by CODEX^5^ gave us the opportunity to use CellCharter to characterize and compare spatial clusters between two conditions: healthy spleen (BALBc model) versus spleen in mice affected by systemic lupus erythematosus (MRL model) (**Fig. 3a**). CellCharter determined stable cluster solutions for n = 4 and n = 11 clusters (**Fig. 3b**), and follow-up analyses were performed on 11 clusters (**Supplementary Figure Fig. 3a**). Cells were assigned to all spatial clusters in all samples (**Fig. 3c**), although, whereas BALBc samples showed a similar distribution of cluster proportions, MRL samples appeared more heterogeneous, particularly among stages of the disease (early, intermediate and late). Using available cell type annotations, CellCharter determined cell type enrichments within each cluster, and consistent with the notion of a cellular niche, most clusters were enriched for specific combinations of cell types (**Fig. 3d** and **Supplementary Figure Fig. 3a**). For example, we identified a B follicle cluster (C3) that became less prevalent with increasing stages of the disease, as opposed to a B-PALS boundary cluster (C2) (**Fig. 3e-f**, insets 1-2-3). Other areas of the spleen were split into multiple clusters based on the prevalence of different cell types. For example, two separate clusters were associated with trabecular structures (C10, C11), one of which was highly enriched in granulocytes, marked by *Ly6G* expression (C11), and expanded with the emergence and progression of the disease (**Fig. 3e-f**, insets 4-5-6). Finally, one cluster was associated with staining artifacts or missing markers (C1) (**Supplementary Figure Fig. 3b**).

**Figure 3:**
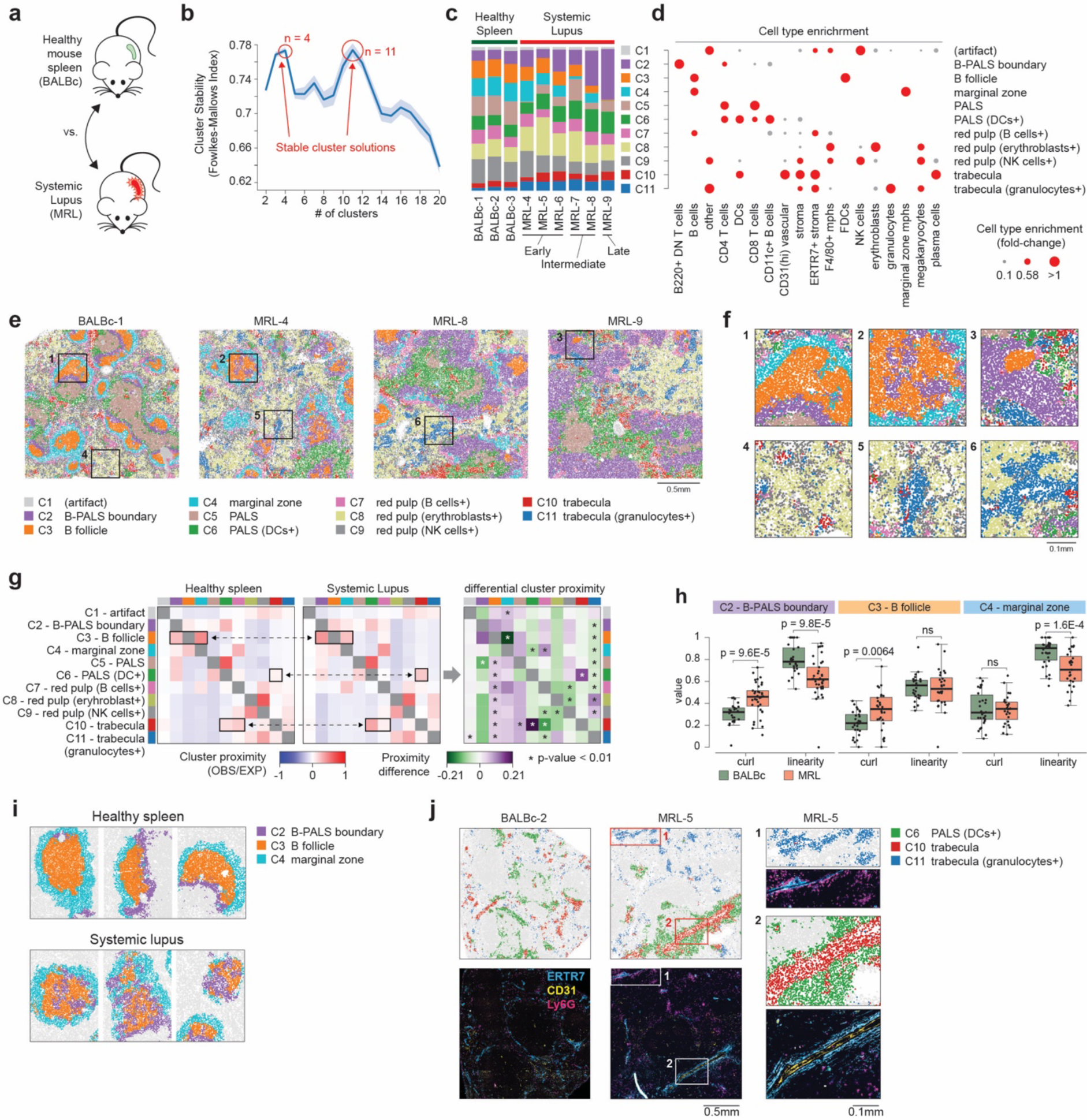
Spatial cellular niches in the spleen of healthy mice and mice affected by systemic lupus. **a)** We applied CellCharter to a spatial proteomics dataset of spleens of healthy mice (BALBc) and mice with systemic lupus erythematosus (MRL). **b)** CellCharter cluster stability (y axis) for a range of number of clusters (x axis). Most stable cluster solutions are highlighted. Data are presented as mean values with a 95% confidence interval. **c)** CellCharter spatial cluster proportions in each sample. MRL samples are grouped based on disease stage (early, intermediate, late). **d)** Cell type enrichment in each spatial cluster. Based on cell type enrichment and spatial location in the tissue, every cluster was associated with an anatomical area of the spleen (annotated on the right). **e)** Spatial clusters in one healthy sample (BALBc-1) and three systemic lupus samples at different disease stages: MRL-4 (early), MRL-8 (intermediate), MRL-9 (late). **f)** Areas 1, 2, and 3 zoom in on representative clusters associated with B follicles. Areas 4, 5, 6 zoom in on representative clusters associated with granulocyte-enriched trabecular structures. **g)** Cluster NE (left and center heatmap) and differential NE (right heatmap) between spatial clusters in healthy mice and in mice affected by systemic lupus. Examples of differentially enriched neighborhoods are highlighted. **h)** Curl and linearity values (y axis) for clusters C2 (left, n = 64), C3 (center, n = 57), and C4 (right, n = 57) in healthy (green) and systemic lupus samples (red). p-values were computed by two-tailed Wilcoxon test, NS, not significant. **i)** Representative examples of C2, C3, and C4 cluster components from healthy (upper) and systemic lupus samples (lower). The boxes show the quartiles of the dataset, whereas the whiskers extend to points that lie within 1.5 interquartile ranges of the lower and upper quartile. **j)** Representative examples of C6, C10, and C11 cluster components from BALBc-2 and MRL-5 samples (upper) and matching immunofluorescence staining images for the indicated markers (lower). Insets 1 and 2 highlight the differential composition of trabecular clusters C10 and C11.

To further appreciate tissue remodeling from normal spleen to systemic lupus, we performed cluster NE and differential cluster NE analysis. Significant NE changes between BALBc and MRL samples concerned the B follicle, marginal zone and B-PALS clusters. In MRL samples, the B follicle cluster (C3) decreased interaction enrichment with the marginal zone (C4) in favor of interactions with the B-PALS cluster (C2) (**Fig. 3g**). Comparison of cluster shapes revealed that both the B-PALS and B follicle clusters significantly increased their curl values, whereas B-PALS and marginal zone clusters significantly lost linearity (**Fig. 3h**). These shape differences indicated increased irregularity of the clusters and loss of the tissue architecture characteristic of normal spleen anatomy (**Fig. 3i**). Overall, changes in cluster proportions (**Fig. 3c**), differential cluster NE (**Fig. 3g**), and shape comparisons (**Fig. 3h**) reflected a gradual expansion and infiltration of T cells from the B-PALS cluster into the B follicle (**Supplementary Figure Fig. 4a**), which is consistent with a germinal center reaction after infection^51^. Differential cluster NE analysis also showed that granulocyte-enriched trabeculae (C11) were restricted in the red pulp enriched in B cells (C7), whereas the other trabecular cluster (C10) was in contact with the PALS cluster, which was enriched in dendritic cells (C6) (**Fig. 3g**). Interestingly, cluster C11 lacked expression of *CD31* (**Supplementary Figure Fig. 4b**), a marker shown to be absent in spleen capillaries within the red pulp but present in the central arteries that are surrounded by PALS^52^. Hence, in MRL samples, CellCharter captured the emergence of two distinct trabecular niches establishing preferential spatial interactions that were not present in the normal spleen: a *CD31*^+^/*Ly6G*^-^ cluster in proximity to PALS and a *CD31*^-^/*Ly6G*^+^ cluster within the red pulp (**Fig. 3j**). Overall, these results demonstrate the potential of CellCharter to characterize remodeling of biological niches in different conditions.

### Using CellCharter to decipher intratumor heterogeneity

Intratumor heterogeneity is characterized by both heterogeneous cancer cell populations, or *cancer cell states*, and diverse composition of the tumor microenvironment (TME)^53–56^. In this context, spatial molecular profiles of multiple tumor samples offer the opportunity to decipher how tumor and TME cell populations organize and interact in the tissue. Here, we used CellCharter to analyze data from 8 non-small cell lung cancer (NSCLC) tissue sections derived from 5 patients: 4 lung adenocarcinoma (LUAD) and 1 lung squamous cell carcinoma (LUSC) (**Fig. 4a**). All tissue samples were previously analyzed using the Nanostring CosMx spatial transcriptomics platform^41^, which assayed mRNA expression for 960 genes at single-cell resolution.

**Figure 4:**
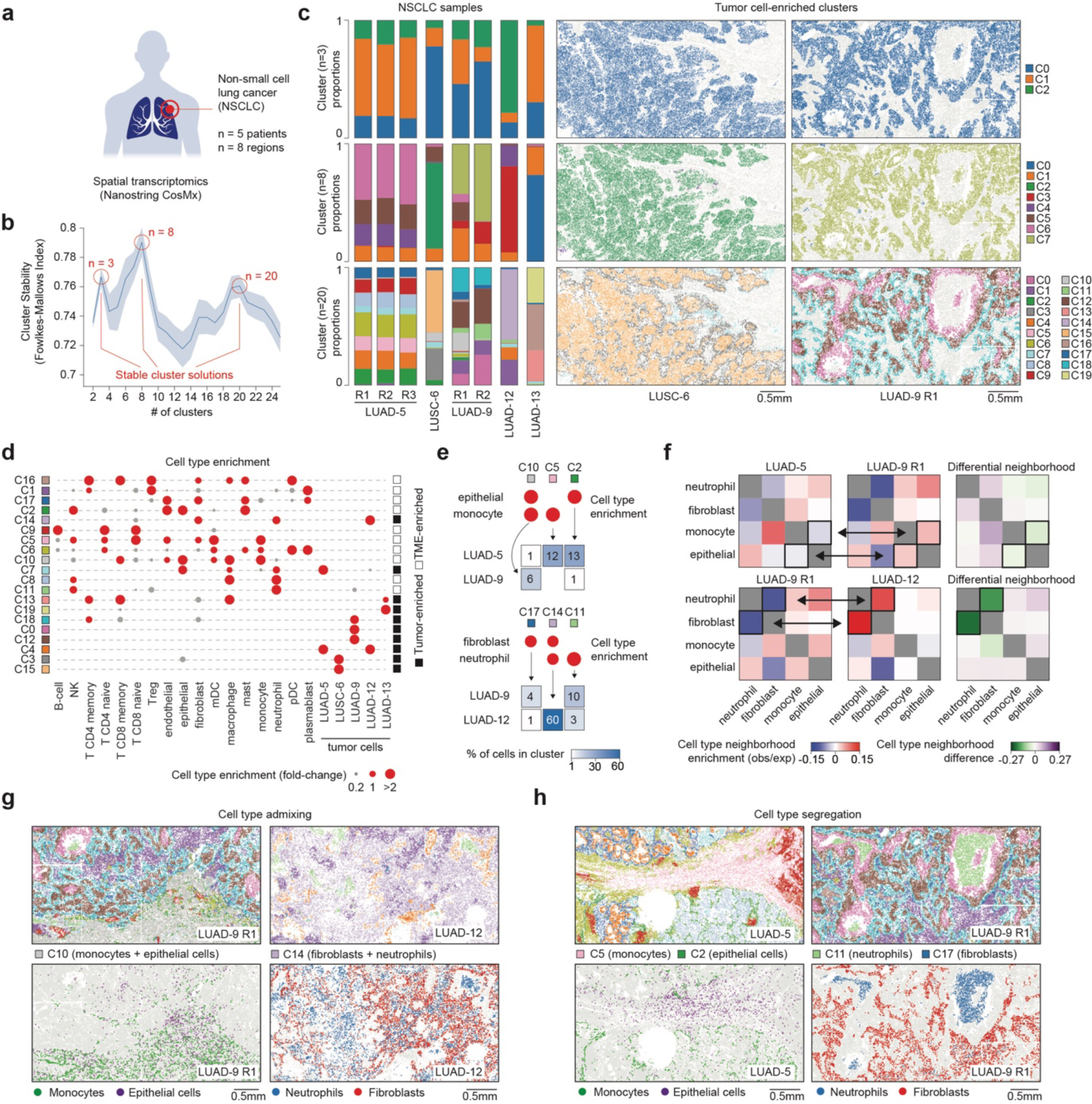
Spatial transcriptomics in non-small cell lung cancer. **a)** CellCharter was applied to a single-cell spatial transcriptomics dataset comprising 8 slides from 5 NSCLC patients. **b)** CellCharter cluster stability (y axis) for a range of numbers of cluster (x axis). Most stable cluster solutions are highlighted. Data are presented as mean values with a 95% confidence interval. **c)** CellCharter spatial cluster proportions in each sample for three numbers of cluster solutions (left) and the corresponding cell labels for tumor cell-enriched clusters in representative samples (right). (LUAD: lung adenocarcinoma, LUSC: lung squamous cell carcinoma.) **d)** Cell type enrichment in each spatial cluster (n = 20 clusters). **e)** Cell type enrichment (red circles) of the indicated cell types (rows) in the indicated spatial clusters (column) and cell type percentages (heatmap) of the indicated tumor sample (rows) in the indicated spatial clusters (column). **f)** Cluster NE (left and center heatmaps) and differential NE (right heatmap) between the indicated cell types in the indicated tumor samples. Examples of differentially enriched neighborhoods are highlighted. **g-h)** Examples of cell type admixing (g) and cell type segregation (h). Spatial cluster assignments (upper) and cell type labeling (lower) for the indicated clusters and cell types in the indicated samples.

CellCharter identified three possible stable solutions with n = 3, 8 or 20 clusters (**Fig. 4b**). Within each solution, CellCharter identified “tumor-enriched” clusters, i.e., clusters that were largely composed of tumor cells (85-90%) and where more than 90% of tumor cells were found (**Supplementary Table 1**). Interestingly, with n = 3 clusters, tumor areas from all 8 samples were largely grouped into one main tumor-enriched cluster that was shared among all patients (**Fig. 4c** - top). With n = 8 clusters, the majority of patients exhibited one private tumor-enriched cluster (**Fig. 4c** - center). Lastly, with n = 20 clusters, each patient exhibited multiple private tumor-enriched clusters, potentially reflecting distinct cancer cell states (**Fig. 4c** – bottom and **Supplementary Figure Fig. 5a**). Hence, to a certain extent, CellCharter stable cluster solutions reflected a hierarchy of biological entities: cancer, individual tumors, and intratumor cell states. To explore spatial features of intratumor heterogeneity, we focused on the 20-cluster solution. Cell type enrichment analysis confirmed the presence of tumor-enriched clusters and clusters characterized by distinct combinations of immune and stromal cell types, referred to as TME-enriched clusters (**Fig. 4d**). Tumor-enriched clusters were almost invariably patient-specific but shared between independent samples derived from the same patient (**Fig. 4c** – barplot). Although TME-enriched clusters were often shared among patients (**Supplementary Figure Fig. 5b**), those that were not often reflected distinct admixing of the same cell types. For example, LUAD-5, LUAD-9 and LUAD-12 exhibited a similar proportion of fibroblasts, neutrophils, monocytes, and normal epithelial cells (**Supplementary Figure Fig. 5c**). However, epithelial cells and monocytes were both enriched in cluster C10 in LUAD-9 but clustered separately in LUAD-5 (C2 and C5) (**Fig. 4e** - top). Similarly, neutrophils and fibroblasts were both enriched in cluster C14 in LUAD-12 but were enriched in different clusters (C11 and C17) in LUAD-9 (**Fig. 4e** - bottom). Cell-type NE analyses confirmed that these clusters were driven by spatial admixing, with epithelial cells and monocytes spatially closer than expected in LUAD-9 but not in LUAD-5, and neutrophils and fibroblasts spatially closer than expected in LUAD-12 but not in LUAD-9 (**Fig. 4f**). Visual inspection of clusters and cell types in the tissues confirmed that immune and stromal cell types that were present in similar proportions in these tumors exhibited distinct spatial organizations, corresponding to either cell type admixing (**Fig. 4g**) or cell type segregation (**Fig. 4h**). These results demonstrate well the added value of including spatial information in single-cell molecular profiles.

Distinct tumor-enriched clusters in the same patient exhibited preferential interactions with distinct TME-enriched clusters (**Fig. 5a**). In patient LUAD-9, we found 2 major tumor-enriched clusters (C0, C12) and a third cluster composed of both tumor cells and CD4^+^ T cells (C18) (**Fig. 4d**). C0 exhibited frequent interactions with a cluster composed of neutrophils and NK cells (C11), whereas C12 mostly interacted with C18 (**Fig. 5b** and **Supplementary Figure Fig. 5c**). Differential expression analysis between clusters C0 and C12 (**Supplementary Table 2**) revealed that upregulated genes in C0 compared with C12 comprised the hypoxia-inducible gene *NDRG1*, which may also promote stem-like phenotypes and epithelial-to-mesenchymal transition (EMT) in lung cancer^57,58^, the angiogenic factor *VEGFA*, which is also induced by hypoxia^59^, and several genes associated with cytokine signaling and neutrophil chemotaxis. Prominent examples were *S100A8* and *S100A9*, and chemokine-encoding genes *CXCL1*, *CXCL2*, and *CXCL3* (**Fig. 5c**), which encode for neutrophil attractants^60^ that have been shown to promote tumor invasion and metastatic capacity^24^. Gene set enrichment analysis confirmed that genes encoding for proteins involved in cytokine signaling and neutrophil chemotaxis were enriched among upregulated genes in the tumor-enriched cluster C0, consistent with the spatial proximity between this cluster and the neutrophil-enriched cluster C11 (**Fig. 5c**). Upregulated genes in C12 compared with C0 were instead enriched for cell proliferation markers, such as *MKI67*, and comprised genes encoding for fibroblast growth factor receptors *FGFR1* and *FGFR2*, and for the histone modifier *EZH2* (**Fig. 5c**), which is frequently overexpressed in aggressive lung adenocarcinoma^61^. Consistent with the upregulated markers in the two clusters, gene signature scores associated with cytokine signaling, response-to-hypoxia, and EMT were higher in cluster C0 than in C12 (**Supplementary Figure Fig. 6a** and **Supplementary Table 3**) and colocalized in the tumor niche surrounding the neutrophil-enriched cluster C11 in independent tumor samples (**Fig. 5d** and **Supplementary Figure Fig. 6b-d**). Conversely, these signatures exhibited an anticorrelated spatial gradient with a cell proliferation signature (**Fig. 5d**, **Supplementary Figure Fig. 6e** and **Supplementary Table 3**). In particular, the response-to-hypoxia signature scores in tumor cells were anticorrelated with the distance between tumor cells and neutrophils that were found in cluster C11 (**Supplementary Figure Fig. 7a**). These results were largely recapitulated using STAGATE, indicating that they are not dependent on the adopted tool (**Supplementary Figure Fig. 7b-d**), although STAGATE failed to discriminate among neutrophils interacting with C0 and other neutrophils in the tumor, favoring cell type features to spatial interactions (**Supplementary Figure Fig. 7e-g**). Overall, the tumor-enriched clusters C0 and C12 effectively represented two cancer cell states coexisting in the same tumor and exhibiting distinct tumor-TME interactions.

**Figure 5:**
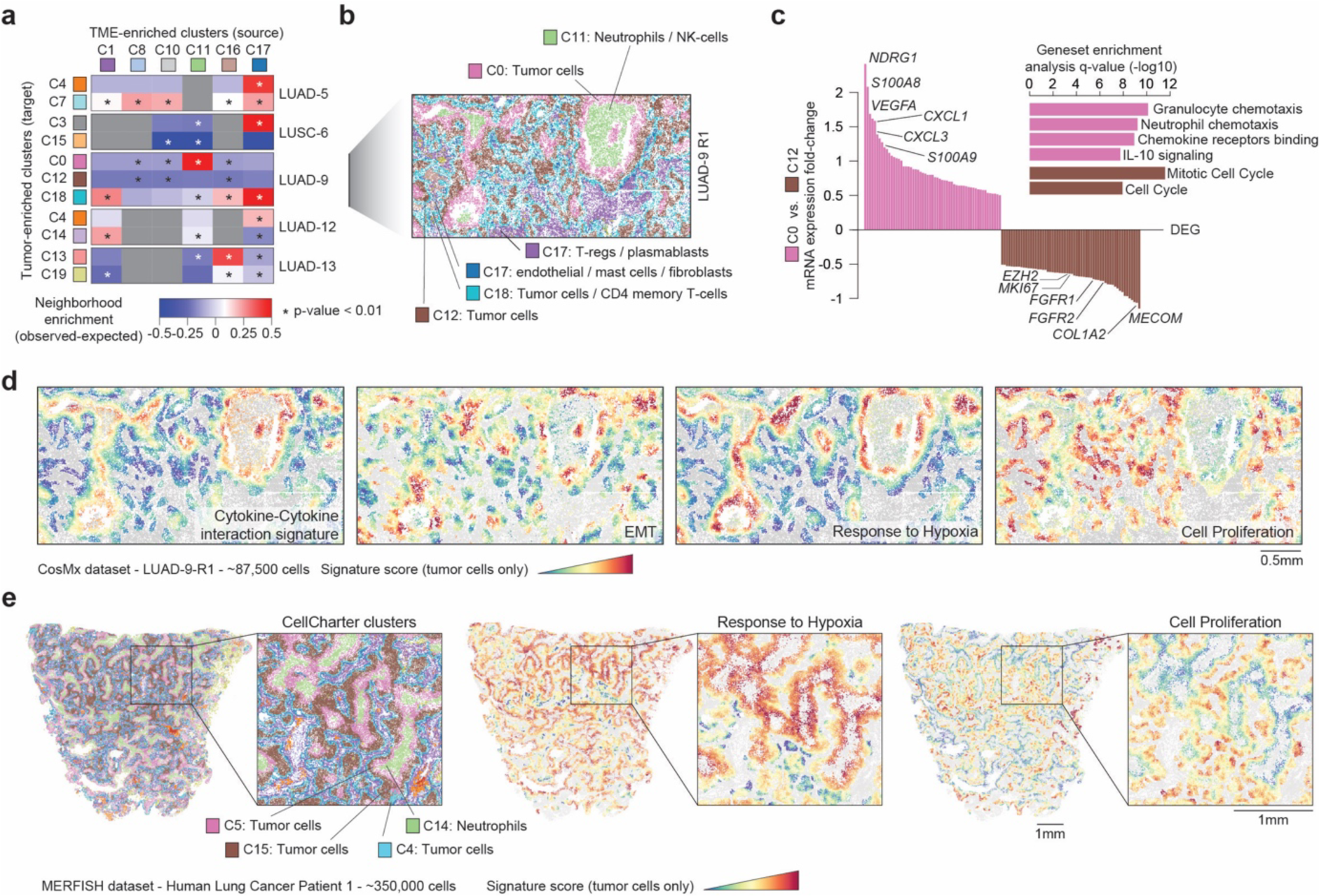
Spatial interactions among cancer cell states and the tumor microenvironment (TME) in lung adenocarcinoma. **a)** NE analysis between the TME-enriched clusters (source) and tumor-enriched clusters (target). Only TME clusters for which there is a significant NE with at least one tumor-enriched cluster are shown. p-values computed by unpaired two-sided t-test. **b)** TME-enriched and tumor-enriched clusters in LUAD-9 R1 (CosMx). **c)** mRNA expression (log2 fold change; y axis) of differentially expressed genes (DEGs; x axis) between the clusters C0 and C12 in patient LUAD-9 (left) and gene set enrichment analysis of genes upregulated in C0 (pink) and C12 (brown) (right). **d)** Gene expression signature scores of tumor cells in LUAD-9 R1 for four different gene signatures. **e)** TME-enriched and tumor-enriched clusters (left) and gene expression signature scores of tumor cells for two gene signatures (center and right) in a LUAD sample analyzed by MERFISH.

To further investigate and validate this lung cancer spatial niche in independent datasets, we collected data from 8 tumor samples that were analyzed by Multiplexed Error-Robust Fluorescence in situ Hybridization (MERFISH by Vizgen; 500 markers), among which there were 2 lung cancer tissues, and a LUAD cohort comprising a tissue microarray of 416 cores analyzed by IMC (35 markers). CellCharter was able to effectively and efficiently jointly analyze all samples in each dataset (**Supplementary Figure Fig. 8a-b**).

In the 2 lung cancer tissue samples analyzed by MERFISH, CellCharter identified n = 20 clusters (**Supplementary Figure Fig. 8c**), out of which 3 tumor-enriched clusters (C4, C5, C15) exhibited similar features to those identified in the CosMx dataset (**Fig. 5e** and **Supplementary Figure Fig. 8d-e**). In particular, cluster C5 was significantly interacting with a neutrophil-enriched cluster (C14) (**Supplementary Figure Fig. 8e**), and exhibited high scores for the response-to-hypoxia gene signature (**Fig. 5e**), which was spatially anticorrelated with a cell proliferation signature. In the IMC dataset (**Fig. 6a**), we used CellCharter to identify n = 30 spatial clusters, which recapitulated the prognostic associations discovered in the original study^24^ (**Supplementary Figure Fig. 8f**). Next, we determined tumor-enriched and TME-enriched spatial clusters and their spatial interactions (**Fig. 6b**). Among the tumor-enriched clusters identified by CellCharter was cluster C23, which exhibited high expression of myeloperoxidase (*MPO*) that is abundant in neutrophils and release of which has been associated with tumor invasion and progression^62^, and hypoxia-inducible factor *HIF1A*, indicative of the activation of a response-to-hypoxia state (**Fig. 6c**). C23 exhibited significant interactions with multiple clusters enriched for neutrophils, most of all cluster C7 (**Supplementary Figure Fig. 8g**). Representative images across multiple patients confirm a spatial niche in which the dense neutrophil-enriched cluster C7 is invariably surrounded by the tumor-enriched cluster C23, which activates a response to hypoxia (**Fig. 6d**). To investigate the prognostic value of this niche, we first analyzed neutrophil-specific gene signatures that discriminate between tumor-associated neutrophils (TANs) and neutrophils associated with normal adjacent tissue (NANs) in NSCLC^63^. In multiple independent lung adenocarcinoma cohorts^11,64–71^, the response-to-hypoxia signature was highly correlated with TAN infiltration but showed no correlation with NAN (**Fig. 6e** and **Supplementary Figure Fig. 9a-b**). Strikingly, in several datasets, multivariate Cox regression analysis found a significant association with worse prognosis for response-to-hypoxia and TAN signatures, but not for NAN (**Fig. 6f**), and the response-to-hypoxia signature retained prognostic value when these signatures were jointly analyzed (**Supplementary Figure Fig. 9c**). Overall, CellCharter was instrumental in revealing a tumor-TME niche across multiple lung cancer patients and cohorts, which was consistently associated with worse patient prognosis.

**Figure 6:**
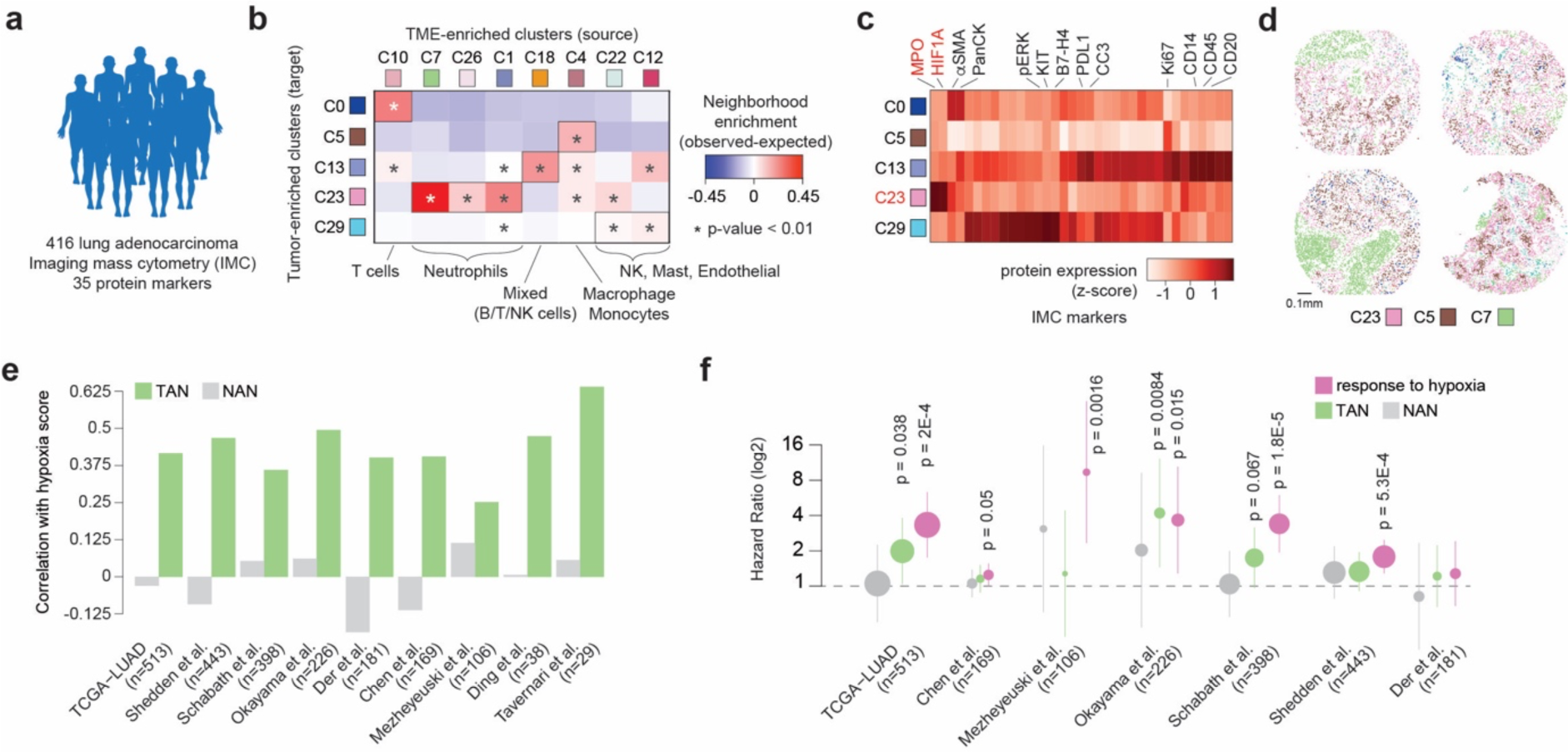
TANs and response-to-hypoxia are associated with worse patient prognosis. **a)** CellCharter was used to investigate 416 LUAD samples analyzed by IMC. **b)** NE analysis between the TME-enriched clusters (source) and tumor-enriched clusters (target) in the IMC dataset. Only TME clusters for which there is a significant NE with at least one tumor-enriched cluster are shown. Cell types enriched in the TME-enriched clusters are annotated. p-values computed by unpaired two-sided t-test. **c)** Normalized protein abundance (Z-score) for the 35 markers analyzed by IMC in the tumor-enriched clusters. Representative markers are annotated at the top. **d)** TME-enriched and tumor-enriched clusters in 4 representative cores of tissue microarray of LUAD samples analyzed by IMC. **e)** Correlation values (Pearson’s coefficient) between response-to-hypoxia signature scores and either TAN signature scores (green bars) or NAN signature scores (gray bars). **f)** Hazard ratios and p-values computed from multivariate Cox regression analysis for TAN (green), NAN (gray) and response-to-hypoxia (pink) gene signature scores. Hazard ratios are presented as mean values with a 95% confidence interval.

## DISCUSSION

The design of computational tools to analyze data from emerging technologies faces two major challenges: 1) anticipating which technologies will ultimately prevail, and 2) anticipating the needs and requirements of analyses that are not yet possible. Currently, spatial molecular technologies vary greatly in throughput and resolution, and spatially resolved cohorts are limited in size and/or number of markers. As technologies mature, multiple layers of spatial omics data will be generated for large collections of samples. We designed CellCharter to be technology-agnostic and highly scalable. We tested CellCharter on different technologies and identified meaningful cell niches in all applications, recapitulating tissue morphologies and biological processes. Importantly, CellCharter was developed with a modular architecture, in which only the dimensionality reduction and batch correction steps depend on the data type, and it is fully compatible with the scverse ecosystem^72^. This design will facilitate the analysis of different spatial data modalities, such as epigenetic features^73^, copy number alterations^74^, chromatin accessibility^75^, and multimodal profiles^17,76,77^. Molecular layers could be combined with cell morphology and tissue histology derived from hematoxylin and eosin (H&E) staining. Deep neural networks trained on large collections of H&E images^78^ together with VAE already used in CellCharter allow to integrate molecular features with tissue histology. Spatial niches determined from this combination could empower digital pathology and serve as prognostic markers in the clinic.

Our results on NSCLC effectively demonstrated the potential of spatial omics profiles to investigate intratumor heterogeneity in terms of both cell-intrinsic features and cell-cell interactions. We discovered a lung cancer cellular niche in which tumor cells characterized by a response-to-hypoxia and EMT transcriptional state surrounded a cluster enriched for TANs. These spatial interactions prompt mechanistic hypotheses regarding the emergence of distinct cancer cell states. Indeed, tumor growth leads to the emergence of hypoxic and necrotic regions, within which cells secrete neutrophil-recruiting chemokines such as *IL-8*, *IL-6*, *CXCL1*, *CXCL2*, *CXCL5*, *CXCL8* and *SOD2*^79–81^, several of which were overexpressed by tumor cells in contact with TANs compared with other tumor cells in the same samples (**Supplementary Table 2**). Neutrophil recruitment can further enhance this signaling and, importantly, promote angiogenesis, cancer cell migration and EMT. Our results hence suggest a positive feedback between these interacting cells promoting cancer cell state transition.

Ultimately, spatial and non-spatial single-cell omics data provide complementary information that together give a holistic definition of *cell state* based on both a cell’s intrinsic molecular features and its set of interactions. Indeed, we showed that the same proportions of the same immune cell populations can exhibit spatial admixing or spatial segregation in different patients. With the extent of immune infiltration and immune cell type composition entering the clinic as markers of therapeutic response^82,83^, it will be interesting to explore the functional and prognostic impact of cell type admixing. Overall, cell spatial coordinates introduce a new critical layer of information to decode phenotypic heterogeneity from molecular data. In this context, CellCharter offers a flexible and scalable solution to interpret spatial information and harness its potential.

## METHODS

### CellCharter: Spatial clustering

Spatial clustering groups spots or cells based on the features of the spot/cell itself, as well as the features of its surrounding neighbors. CellCharter’s approach to spatial clustering involves several steps. Firstly, it constructs a network using the coordinates of the spots/cells. Then, it applies dimensionality reduction and batch effect removal. Next, for each cell, it aggregates its features with those of its neighboring cells. Finally, clustering is performed on the aggregated matrix.

#### Spatial network construction

We represent spatial omics data as networks, where spatial locations are represented as nodes. These nodes are connected by edges if they are in close proximity to each other. Depending on the technology used, we employed different approaches implemented in the Squidpy library^46^ (v1.3.0) to construct the network. For the Visium and RNA+ATAC data, which exhibit a regular structure, we assigned the 6 and 4 closest surrounding spots, respectively, as neighbors for each spot. For the CODEX, CosMx, MERFISH, and IMC data, we constructed the network using Delaunay triangulation^84^. However, Delaunay triangulation can result in long edges between cells, especially at the slide borders. Therefore, we eliminated edges between nodes if the distance between them exceeded the 99th percentile of all distances between connected nodes.

#### Dimensionality reduction and batch effect removal

For dimensionality reduction and batch effect removal, we used variational autoencoders (VAEs) with omics-specific priors. VAEs are a type of neural network that learns low-dimensional embeddings capable of accurately reconstructing the original input data. They have emerged as valuable tools for modeling transcriptomics, epigenomics, proteomics, and multi-omics data, largely due to the incorporation of omics-specific priors. The flexibility in customizing these prior distributions enables VAEs to adapt to the unique data distribution inherent to each omics assay. Furthermore, the scalability of VAEs makes them particularly suited for modeling large-scale datasets.

Spatial transcriptomics simultaneously measures thousands of genes, resulting in high-dimensional data. We used scVI^43^ to perform dimensionality reduction and batch effect removal of the spatial transcriptomics and spatial epigenomics samples (Visium, CosMx, MERFISH, and RNA+ATAC). The transcriptomics data was modeled using a zero-inflated negative binomial distribution^43^, while the chromatin accessibility data was modeled with a Poisson distribution^85^. Spatial proteomics, which quantifies the expression of tens or hundreds of proteins, has lower dimensionality, making dimensionality reduction potentially unnecessary. However, we applied it using scArches^44^ to decrease noise and, by reducing the number of features, the runtime of clustering in the CODEX samples.

#### Neighborhood aggregation

Incorporating the features of neighboring nodes into a given spot or cell is achieved through *neighborhood aggregation*. This technique involves concatenating the features of a spot/cell with those aggregated from its neighbors. This aggregation spans increasing layers from the considered spot/cell, extending up to a certain layer *L*. Aggregation functions are used to derive a single feature vector from the vectors of multiple neighbors. Commonly adopted aggregation functions include the *mean, standard deviation, minimum*, and *maximum*. Depending on the nature of the relationships we aim to capture with the neighbors, new aggregation functions can be defined. For instance, *standard deviation* can measure the variability of cell phenotypes surrounding a particular spot/cell. Additionally, multiple aggregation functions can be used simultaneously to capture various types of relationships, resulting in a total of *J* aggregations. Let’s denote the matrix of spot/cell features (whether dimensionality-reduced or not) as *X*, *A*^(*l*)^ for *l* ∈ {0, …, *L*} as the adjacency matrix of the neighbors at layer *l*, *f_j_* for *j* ∈ {0, …, *J*} as the aggregation functions, (*X*, *Y*) as the matrix concatenation operation, and *X* ⋅ *Y* as matrix inner product operation. The output of the neighborhood aggregation step is a matrix *Z*:

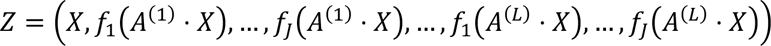

#### Clustering

The matrix *Z* contains information about the spots/cells and their neighbors. As a result, the final step involves performing clustering on *Z* using a Gaussian Mixture Model (GMM). CellCharter uses an implementation of GMMs from the [PyCave](https://github.com/borchero/pycave) library (v3.2.1), which is optimized to run efficiently on either CPUs or GPUs for multiple samples simultaneously using PyTorch (v1.12.1). We chose Gaussian Mixture Models (GMMs) as our clustering method because they can form clusters of varying sizes and distributions and handle diverse feature scales. SUch flexibility allows the concurrent use of various data sources, such as multi-omics data.

#### Cluster stability

Clustering with a GMM requires specifying the desired number of clusters *K*. Therefore, CellCharter offers a procedure based on the stability of the clustering to identify the optimal candidates for *K*. The goal is to identify the values of *K* that yield consistent clustering results across multiple runs with *K*, *K* − 1, and *K* + 1 clusters. The user specifies the range of values of *K* to scan and the maximum number of repeated runs *R* to perform for each value of *K*. For each *K* within the specified range, CellCharter executes a single clustering run with *K* clusters. Following this set of clusterings, the average of the Fowlkes-Mallows Index (FMI)^45^ is calculated between the clusters at *K* − 1 and *K*, and between *K* and *K* + 1. The FMI is defined as the geometric mean of *precision* and *recall* between two clustering assignments. Consequently, a higher similarity between clustering solutions at consecutive numbers of clusters corresponds to a higher average FMI. Subsequently, a new set of clusterings is executed, and the average FMI is computed for all combinations of results between the *r* = 2 runs at *K* − 1 and *K*, and at *K* and *K* + 1. This procedure is repeated until the *r* = *R*. To enhance computational efficiency, it is possible to set that, for each new additional run *r*, if the Mean Average Percentage Error between the average FMIs at *r* − 1 and *r* falls below a user-defined *tolerance* value, then the process is deemed to have converged and completes without needing to reach *r* = *R*. Since the process generates up to *R* models for every candidate number of clusters, once a value for *K* is determined, CellCharter selects the model with the highest marginal likelihood out of the *r* models.

### CellCharter: Cell type enrichment

Identifying the cell type composition of spatial clusters can facilitate their characterization. We can compute the enrichment of cell type *t* in cluster *c* as the ratio between the observed and the expected proportions of cells of type *t* in *c*. One possible approach to estimate the expected value involves computing the average proportion from multiple permutations of the cell type labels. This method also allows to estimate an empirical p-value, defined as the frequency at which the observed proportion exceeds the expected proportion. Alternatively, under the assumption that, given a large number of permutations, the expected proportion converges to the proportion of cells of type *t* across all samples, we can derive an analytical formulation for the expected value. While this approach is more efficient, it doesn’t allow for significance estimation. Finally, the enrichment values are then log_2_-normalized. This results in a zero value if the likelihood of *t* being in *c* is equivalent to random chance, a negative value for depletion, and a positive value for enrichment. In the CODEX mouse spleen dataset, a cell type was considered enriched with a log_2_ fold-change greater than 0.58, equivalent to a fold-change of 1.5. In the CosMx NSCLC, MERFISH, and IMC lung cancer datasets, a cell type was considered enriched with a log_2_ fold-change greater than 1, equivalent to a fold-change of 2.

### CellCharter: Neighborhood enrichment

#### Analytical neighborhood enrichment

Given cells partitioned into *K* groups *C*_*i*_ for *i* ∈ {1, …, *K*} and an adjacency matrix *A* of the spatial network composed of *V* nodes and *E* edges, let *k*_*v*_ denote the node degree of cell *v*. The proximity between two groups can be computed from their neighborhood enrichment, which measures the likelihood of cells from two groups to be connected by an edge, compared to random chance. We developed an analytical formulation for the neighborhood enrichment between groups *C*_*i*_ and *C_j_*, which is computed as the ratio between the observed and the expected number of links connecting cells in *C*_*i*_ and cells in *C_j_*.

Formally, the observed value between two clusters is the total number of links between them:

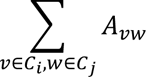

The expected value is obtained from the probability of two nodes in groups *C*_*i*_ and *C_j_*to be connected, which is proportional to the product of their degrees. The expected number of links between *C*_*i*_ and *C_j_* is then the sum of the expected number of links between each pair of nodes where one node is in *C*_*i*_ and the other is in *C_j_*:

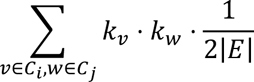

The neighborhood enrichment calculated through permutation converges to the analytical one, producing a matrix of size *K* × *K*. Although the analytical version is considerably more efficient, it does not allow for the estimation of a p-value for a specific neighborhood enrichment.

#### Asymmetric neighborhood enrichment

The symmetric formulation of neighborhood enrichment fails to capture unbalanced connectivities between groups. Therefore, we developed an asymmetric version of neighborhood enrichment that considers the proportion, rather than the number, of edges from cells in *C*_*i*_ connected to cells in *C_j_*. In this setting, the neighborhood enrichment between *C*_*i*_ and *C_j_* may differ from the neighborhood enrichment between *C_j_* and *C*_*i*_. The observed value between groups *C*_*i*_ and *C_j_* is the number of edges between *C*_*i*_ and *C_j_* divided by the total number of edges of the cells in group *C*_*i*_:

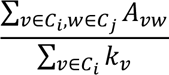

The expected value is the proportion of edges connected to *C_j_*:

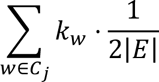

In this context, the asymmetric neighborhood enrichment was computed as the difference between the observed and expected values, rather than as a ratio.

#### Differential neighborhood enrichment

The differential neighborhood enrichment between two conditions is defined as the difference between the neighborhood enrichment matrices for those conditions. If the user needs to estimate the significance of differences in neighborhood enrichment, an associated p-value can be calculated through random sampling (with replacement) of the condition label for each sample. The p-value for the differential neighborhood enrichment between groups *C*_*i*_ and *C_j_*is the proportion of cases where the permuted value is greater than the observed value, when the latter is positive, and the permuted value is lower than the observed value, when the latter is negative. Estimating p-values is optional and requires computing the neighborhood enrichment several times. However, the development of an analytical version of the non-differential neighborhood enrichment makes this process computationally feasible.

### CellCharter: Shape characterization

The same cellular niche can be present in multiple tissues or even in various locations within a single tissue sample. This suggests the existence of multiple cluster components, which are connected components of cells belonging to the same spatial cluster. To characterize the shape of a spatial cluster, it is necessary to identify its cluster components, determine their boundaries, and compute the value of the metrics that we developed to describe shape (linear, round, circular, and irregular). For determining the boundaries of a cluster component, we developed a new technique based on alpha shapes^86^. For each component, we computed the alpha shape with a starting value of alpha that depends on the resolution of the data. If the alpha value was too small, the alpha shape of the component would yield multiple separated polygons. In such cases, we doubled the alpha value and recomputed the alpha shape. This procedure is iterated until an alpha shape composed of a single polygon is obtained. Ultimately, the boundaries of the cluster component are defined as the alpha shape with the minimum alpha value that results in a single polygon. To keep shapes simple, holes with an area relative to the boundary area smaller than a threshold *a* (e.g., 0.1) were removed. Consequently, a boundary can contain holes with an area greater than *a* times the area of the boundary. Once we determined the boundary of each component, we used the [Shapely](https://github.com/shapely/shapely) library to compute geometric information such as its perimeter *P*, area *A*, its minor and major axes from its minimum rotated rectangle, which are used to compute the following four shape metrics.

#### Curl

Curl measures how twisted or curved is a cluster. It is computed as one minus the ratio between the major axis of the minimum rotated rectangle that fits the polygon and the fiber length. Circular and irregular clusters exhibit a high curl.

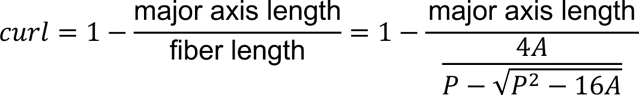

#### Elongation

Upon fitting the minimum rectangle surrounding a cluster, a linear cluster will result in a rectangle with the minor edge significantly shorter than the major edge. Elongation is defined as one minus the ratio between the lengths of the minor and major axes.

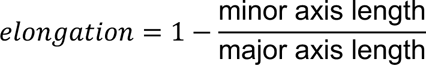

#### Linearity

We developed a new metric termed linearity, which measures the ability of a cluster component to approximate a linear path. Initially, we computed the skeleton of the polygon using a type of skeletonization^87^ implemented in the scikit-image library^88^. Subsequently, we used the [sknw](https://github.com/Image-Py/sknw) library to obtain a weighted network, wherein a node is a juncture between lines of the skeleton, and two nodes are linked by an edge if they are connected by a line in the skeleton. The weight of an edge is the length of the line connecting the two junctures. Linear and circular clusters tend to have a skeleton that is composed of a single main axis, with a few short lines branching out, while the skeleton of round and irregular clusters has numerous bifurcations of similar length. Thus, linearity is defined as the length of the longest path in the network divided by the total length of the network. To also account for circular clusters, we include as paths all the possible cycles which form a basis for cycles in the network, computed using the NetworkX library^84^.

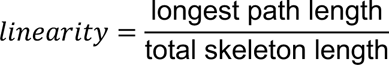

#### Purity

Cells of a cluster may be locally intermixed with cells of other clusters. If the intermixing does not exceed the level at which the cells of the cluster form a connected component, cells from different clusters could be found within the boundaries of a cluster component.

Purity measures the degree of cluster intermixing within a component. Very compact clusters will exhibit higher purity than sparse and irregular clusters. Thus, given *N* cells within the borders of a cluster component, for each cluster *k* ∈ {1, …, *K*}, *N*_*k*_ cells are assigned to cluster *k* such that ∑_*k*∈{1,…,*K*}_ *N*_*k*_ = *N*, then the purity of a cluster component of cluster *k*] is defined as:

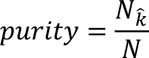

### Benchmarking comparison of CellCharter with existing tools

We compared CellCharter against six established methods for spatial clustering: DR-SC (v.2.7), UTAG (commit 0d3d420), SOTIP (commit d3b762c), SEDR (commit 18616df), BayesSpace (v.1.4.1), and STAGATE (v1.0.0). Since UTAG uses Leiden clustering, the number of clusters can only be controlled through the resolution parameter. Consequently, we modified the clustering procedure to directly select the number of clusters. We executed Leiden clustering starting with a resolution of 0.1. If the obtained number of clusters was lower than the desired number, the process was repeated, increasing the resolution by 0.1. If a clustering solution returned a number of clusters greater, but not equal, to the desired number, we iteratively merged the pairs of clusters with the lowest Euclidean distance until the target number of clusters was reached. The dataset comprises annotated samples of six-layered human dorsolateral prefrontal cortex, processed using the 10x Genomics Visium platform^47^. The samples were divided into two pairs of adjacent slides from each of the three donors, for a total of 12 slides. Almost every Visium spot was assigned to one of the six cortical layers (L1-L6) or to white matter (WM), resulting in a total of seven labels, with a small number of spots for which a label couldn’t be assigned. We evaluated the methods in two settings:

- individual: each sample is clustered and evaluated individually
- joint: all the samples are clustered jointly and evaluated both individually and jointly.

Initially, we divided the 12 samples into a tuning set (samples 151507, 151672, and 151673, selected randomly from each donor) used for hyperparameter tuning, and a test set composed of the remaining 9 samples. Once the hyperparameters were selected, we ran 10 repetitions of spatial clustering for each individual sample and compared the predicted labels of the method against the annotation labels using the Adjusted Rand Index (ARI). For the joint benchmarking, we integrated all 9 test samples, executed 10 repetitions of joint clustering on the concatenated dataset, and assessed the accordance of the predicted labels against the manual annotations by computing the ARI for each sample separately to assess the variability of the results between different samples. Spots with unassigned manual labels were included in the clustering but excluded from the evaluation. Additionally, we measured the running time and total memory (i.e., CPU and GPU combined) requirements for 5 repetitions of joint clustering for each increasing number of samples, from 2 to 12. The benchmarking was conducted on 4 cores of an AMD Epyc2 7402 CPU and, for GPU-compatible methods, an NVIDIA A100 GPU with a memory size of 40GB.

#### Hyperparameter tuning

For each tuning sample, we conducted a grid search, running 5 repetitions for every combination of hyperparameters. The values associated with the maximum mean Adjusted Rand Index (ARI) across samples on the tuning set were selected.

The hyperparameter values candidates and the identified ones are the following:

##### DR-SC

- Highly variable genes (HVGs) or spatially variable genes (SVGs): SVGs
- N. spatially variable genes [500, 1000, 2000, 5000]: 2000

##### UTAG

- N. highly variable genes [500, 1000, 2000, 5000]: 1000
- N. principal components [5, 10, 15]: 10

##### SOTIP

- N. highly variable genes [1000, 2000, 5000]: 2000
- N. principal components [50, 100]: 50
- N. PCA neighbors [15, 30, 50]: 30
- N. microenvironment (ME) neighbors [10, 30, 50]: 50
- N. EMD neighbors [100, 300, 500]: 500

##### SEDR

- N. hidden features of last fully connected layer [10, 20, 30]: 10
- N. hidden features of last GCN layer [4, 8, 16]: 16
- N. principal components [100, 200, 300]: 100

##### BayesSpace

- N. highly variable genes [500, 1000, 2000, 5000]: 1000
- N. iterations [2000, 5000]: 5000
- N. principal components [5, 7, 10, 15, 20, 25, 30]: 15
- γ [1, 2, 3, 4]: 3

##### STAGATE

- N. highly variable genes [1000, 2000, 5000]: 5000
- N. hidden features [256, 512, 1024]: 1024
- Latent size [5, 10, 30, 50]: 30

##### CellCharter

- N. highly variable genes [500, 1000, 2000, 5000]: 5000
- Latent size [5, 10, 15]: 5
- N. neighborhood layers [2,3,4]: 4

#### Joint benchmarking

Joint clustering requires the integration of multiple samples into a single dataset. Given the presence of batch effects between patients, all methods for spatial clustering were preceded by a dimensionality reduction and batch effect removal step. CellCharter already uses scVI to perform dimensionality reduction of the spatial transcriptomics data, which has recently been demonstrated to be among the best data integration methods for transcriptomics data^89^. Unless specified in the original work or documentation, we applied the same batch effect corrections for the same computational environment: Harmony^90^ with default parameters for the R-based methods (BayesSpace); scVI with default parameters and early stopping for the Python-based methods (SEDR, STAGATE, and CellCharter). UTAG specifies ComBat^91^ as the preferred method for batch effect correction. The R-based package DR-SC does not accept dimensionality-reduced data as input since it implements its own dimensionality-reduction technique. Therefore, we used DR-SC without batch effect removal to evaluate the ARI on the ground truth labels but excluded it from the runtime and memory evaluation.

Finally, we compared CellCharter and STAGATE at various degrees of downsampling of the read counts. We used scanpy’s function *pp.downsample_counts* (v1.9.1) to reduce the total number of read counts to 75%, 50%, 25%, 10%, and 1% of the original number. Then, we trained CellCharter (including scVI) and STAGATE on the test set for each downsampled value, using the same hyperparameters found in the original tuning set, and compared the predicted labels against the manually annotated labels using the Adjusted Rand Index (ARI) metric.

#### Evaluation of batch effect correction

To demonstrate the need for batch effect correction, we ran CellCharter using embeddings generated by scVI, maintaining the same hyperparameters as in the joint clustering but omitting the Visium slide as a batch key. Since all samples originate from the same brain region, all spatial clusters should be shared among donors. However, in the absence of batch effect correction, most clusters were unique to the donor (i.e., Visium slide). Incorporating the batch key resulted in clusters that were shared by all three donors.

#### Evaluation of clustering stability

First, we assessed whether 10 were sufficient to achieve convergence for the cluster stability analysis. We ran cluster stability on the DLPFC dataset with the same settings as in the joint clustering, but increased to 25 iterations, and compared the FMI curve with the run at 10 iterations. Additionally, we calculated the change in the FMI curve at each new iteration as the Mean Average Percentage Error between the FMI curves at consecutive iterations.

We compared CellCharter’s cluster stability against other methods to identify the optimal number of clusters, such as the Akaike Information Criterion (AIC), Bayesian Information Criterion (BIC), and Negative Log-Likelihood (NLL). Using the same settings as in the joint clustering, we ran CellCharter’s clustering stability on all 12 samples of the DLPFC dataset, exploring a range of clusters n from 2 to 20 and conducting 10 repetitions. We observed a peak at n = 9. Then, we ran CellCharter at an increasing number of clusters and measured AIC, BIC, and NLL for each n between 2 and 100. BIC showed a global minimum at n=19. Although neither a minimum nor a clear elbow could be identified for AIC and NLL, the number of suggested clusters using these criteria could be selected in the range of n = 15-25.

We tested the consistency in the number of clusters with increasing dataset sizes using a recently published extended version of the pilot DLPFC dataset^92^. It is composed of 30 additional DLPFC samples analyzed with the 10x Visium technology. We thus merged the pilot and extended datasets and ran CellCharter’s cluster stability analysis, using scVI with the study (pilot or extended) as the batch key and the Visium slide as a categorical covariate. The FMI curve for the merged dataset still exhibitedew a peak at n = 9 and additional peak at n = 14 (**Fig. 1h**). Upon computing AIC, BIC, and NLL on the merged dataset for a range between 2 and 100 number of clusters, BIC displayed a global minimum at n = 65. While neither a minimum nor a clear elbow could be identified for AIC and NLL, the number of suggested clusters using these criteria could be selected in the range of n = 30-40.

#### Spatial clustering of multi-omics mouse brain data

We downloaded the Mouse P22 spatial RNA+ATAC dataset and independently ran dimensionality reduction on each modality using scvi-tools^93^. For ATAC-seq, we selected the fragments expressed in at least 1% of the spots and executed scVI with n_hidden = 334 and n_latent = 18, determined automatically as the square of the number of unique fragments and the square of n_hidden, respectively, n_layers = 1 and employed Poisson likelihood as it was shown to outperform binary approaches^85^. For RNA-seq, we performed CPM and log_2_ normalization, selected the 5000 most highly variable genes, and ran scVI with n_hidden = 334, n_latent = 18 (to maintain the same embedding size across modalities), and n_layers = 2. We used the concatenation of the ATAC and RNA embeddings as features for CellCharter’s cluster stability analysis, using a L-neighborhood = 4, a range for the number of clusters between n = 2 and n = 15, and 10 repetitions. We selected the highest peak at n = 10, which was used to cluster the ATAC+RNA data but also the single modality embeddings individually, and compared the results.

### Spatial clustering of CODEX mouse spleen data

We applied CellCharter to a publicly available dataset of mouse spleens, imaged using the CODEX spatial proteomics technology^5^. The dataset compises 3 samples from healthy mice and 6 samples from mice at different stages of systemic lupus erythematosus (SLE), with 30 proteins measured across over 700,000 cells. We used as cell type annotations the ones provided by the authors. First, we removed from the dataset the cells labeled as “dirt” and excluded MHCII from the markers because of inconsistent staining between the two conditions. As the last step of the preprocessing, we computed the network of spatial neighbors for each sample. Then, for each sample, we applied z-score normalization to the markers individually and employed scArches^44^ (v0.5.9) for dimensionality reduction using the trVAE model. We removed the last ReLU layer of the neural network to allow for continuous and real output values. The trVAE model was trained on the dataset using the mean squared error (MSE) loss, two hidden layers of size 128, no MMD, early stopping patience of 5 epochs, and a latent size of 10. We used the latent embeddings of all cells extracted from scArches as features and estimated the optimal number of clusters using our stability analysis with an L-neighborhood of 3, a range for the number of clusters between n=2 and n=20, and 10 repetitions. n=11 and n=4 were the numbers of clusters with the highest Fowlkes-Mallows Index, and we chose n=11 to gain a more fine-grained view of the spatial architecture of the tissues. The model at n=11 clusters with the highest marginal likelihood was used for the labeling of the cells, even though we didn’t find striking differences in the labeling between different runs, given the high Fowlkes-Mallows Index (0.78) at 11 clusters. Then, each cluster was labeled based on its cell type enrichment and the location of its cells in the tissue.

#### Comparison between CellCharter and STAGATE

We compared the running time and label assignment on the CODEX mouse spleen dataset between CellCharter and STAGATE, the two best-performing methods according to our benchmark. For STAGATE, given the large number of cells in the CODEX mouse spleen dataset, fitting the entire dataset into the GPU memory was not feasible. We relied on the batch training strategy, dividing each sample into different subgraphs based on the x and y coordinates and using a subgraph as a batch in the training process. We split every sample into 24 subgraphs, 4 subgraphs based on the x coordinate and 6 subgraphs based on the y coordinate. Given the 9 samples, the split resulted in 216 subgraphs. The spatial network was generated using a radius cutoff of 9000 pixels. We trained STAGATE using a single hidden layer with default parameters: size = 512, latent size = 10, learning rate = 0.001, weight decay = 0.0001, number of epochs = 1000, and performed clustering at 11 clusters. On the other hand, CellCharter did not require any sample splitting. Ultimately, we compared the running time and label assignment of STAGATE against a single run of CellCharter. To assess the quality of the labels, given the absence of ground truth regarding the anatomical area of the cells, we relied on the pathologist annotations^5^ for a visual comparison.

#### Cluster neighborhood enrichment of CODEX mouse spleen data

We computed the cluster neighborhood enrichment for each condition independently, omitting the intra-cluster links to emphasize only interactions between different clusters. We then conducted differential neighborhood enrichment between healthy and systemic lupus samples. We ran 1000 permutations to estimate the significance using a threshold for the p-value of 0.01. Notably, all significant pairs of clusters retained their significance upon Benjamini-Hochberg correction, with an adjusted p-value threshold of 0.05.

#### Shape characterization of CODEX mouse spleen data

For the CODEX dataset, we identified the cluster components for each spatial cluster by isolating the connected components comprising more than 250 cells. Then, for each cluster component, we computed the alpha shape with a starting value of alpha equal to 2000 pixels and a minimum hole area ratio *a* of 0.1.

### Spatial clustering of CosMx NSCLC data

We applied CellCharter to a publicly available dataset of Non-Small Cell Lung cancer (NSCLC) samples derived from the Nanostring image-based spatial transcriptomics CosMx technology^41^. The dataset encompasses 8 samples from 4 lung adenocarcinoma and 1 lung squamous cell carcinoma patients. Three slides from patient LUAD-5 were obtained from serial sections, while the two slides from patient LUAD-9 were obtained from non-serial sections. The dataset comprises 960 genes measured on more than 750,000 cells. We filtered out genes expressed in fewer than 3 cells and cells with fewer than 3 genes expressed, then performed CPM and log_2_ normalization. Subsequently, we ran dimensionality reduction and batch effect removal using scVI from scvi-tools (v0.20.3). We used the default parameters (one hidden layer of size 128, latent size 10, early stopping with patience of 45 epochs), batch effect removal with the sample as batch key, and the patient as a categorical covariate. We used the latent embeddings of all cells extracted from scVI as features, an L-neighborhood of 3, and estimated the optimal candidates for the number of clusters using our stability analysis. This analysis, involving 10 repetitions and a range for the number of clusters from n = 2 to n = 25, suggested n = 3, n = 8, and n = 20 as the best candidates. We selected n = 20 for downstream analysis to highlight cellular niches containing different tumor subpopulations in the same sample.

#### Comparison between CellCharter and STAGATE

For the comparison with STAGATE, we used the same embeddings derived from scVI and used batch training, that splits the samples into multiple subsamples, to be able to fit the samples in memory. Thus, we divided each sample into 25 subgraphs, achieved by splitting based on both the x and y coordinates into 5 sections each, totaling 200 subgraphs. The spatial network was constructed using a radius cutoff of 100 pixels. We trained STAGATE using default parameters and number of clusters n = 20, the same used for CellCharter. For each pair of clusters *i* and *j* we computed the proportion of cells of CellCharter’s cluster *i* that are in STAGATE’s cluster *j*. Cancer cell signature scoring and cell type enrichment were run under the same settings as those in CellCharter.

#### Cluster and cell type neighborhood enrichment of CosMx NSCLC data

Cluster neighborhood enrichment was conducted across all 8 samples together, utilizing CellCharter’s 20 spatial clusters, with p-value estimation performed through 1000 permutations. On the other hand, cell type neighborhood enrichment was executed individually for each sample. In both scenarios, we removed intra-cluster (or intra-cell type) links to highlight only interactions between different clusters (or cell types).

#### Differential expression analysis of CosMx NSCLC data

We used Seurat^94^ to log-normalize the counts from the original, unnormalized data, using a scale factor equal to 10,000. We then used MAST^95^ (v1.24.1) to identify genes differentially expressed between the tumor cells withn the spatial clusters C0 and C12. We selected the genes with a log_2_ fold-change higher than 0.5 and a p-value below 0.05. We performed gene set enrichment analysis of the differentially expressed genes using the GSEA tool^96^ (v.4.2.3). Given that the CosMx NSCLC dataset contains gene expression measurements for 960 genes, we used the pre-ranked version, to avoid a possible bias because of the different gene universe. This approach conducts gene set enrichment analysis against the full list of 960 genes, ranked from the highest to the lowest fold-change. We ran GSEA against the GO:BP (Gene Ontology Biological Process) and the CP:KEGG (Canonical Pathways KEGG) gene set databases at version 7.5.1.

#### Cancer cell signature scoring

We visualized the spatial distribution of tumor cells in LUAD-9 expressing different levels of specific signatures. These signatures were formulated by selecting the most significant GO:BP and CP:KEGG gene sets derived from the differential expression analysis between clusters C0 and C12. Additionally, we included signatures from 41 meta-programs, identified from scRNA-seq data across 24 tumor types^97^ (**Suppl. Table 3)** and we constructed the signature “cell proliferation” by merging the 4 cell-cycle-related signatures. Given a gene set, we computed its score for each cell using the *tl.score_genes* function from scanpy^98^ (v1.9.1). Given the 960 genes measured by the CosMx technology, signatures had a relatively small size (**Suppl. Table 3**). To mitigate the noise caused by the small size of the signatures, we smoothed the signature score for each cell by averaging over the 50 nearest cancer cell neighbors.

#### Cancer cell signature score dependence on neutrophil distance

We assessed the level of the hypoxia signature score of tumor cells at increasing distance from the neutrophils. Specifically, we computed the *L*-hop neighbors for all neutrophils in patients LUAD-9 and LUAD-12, exploring *L* values from 2 to 40. For each *L*-hop adjacency matrix, we isolated links between neutrophils and tumor cells and used CellCharter’s neighborhood aggregation on the hypoxia score to obtain the mean hypoxia score of the tumor cells at distance *L* from the neutrophils. We executed the procedure for all neutrophils, for the neutrophils in the spatial cluster C11, and for the neutrophils not in C11.

### Spatial clustering of MERFISH Human Immuno-oncology data

We assessed CellCharter’s scalability in terms of time and memory using the full MERFISH dataset from Vizgen, composed of gene expression data from 500 genes, spanning 8 tumor types, 16 samples, for a total of more than 8.5 million cells. After filtering out genes expressed in fewer than 10 cells and cells with fewer than 10 genes expressed, we performed CPM and log_2_ normalization. Then, we ran dimensionality reduction and batch effect removal using scVI from scvi-tools (v0.20.3). We used the default parameters (one hidden layer of size 128, latent size 10, early stopping with patience of 45 epochs), and the sample as the batch key. We used the latent embeddings of all cells extracted from scVI as features, an L-neighborhood of 3, and estimated the optimal candidates for the number of clusters through our stability analysis with 10 repetitions and a range for the number of clusters from 2 to 40. To test the memory and time performance for a high number of clusters, we selected n = 29 as it was the highest value preceding a dip in FMI that persisted at least until n = 40. Finally, we ran a single execution of CellCharter with the same hyperparameters and measured the required time and memory.

#### Reference mapping and cell type identification of lung cancer samples

We focused on the two lung cancer samples HumanLungCancerPatient1 and HumanLungCancerPatient2. We used scANVI^99^ from scvi-tools^93^ (v0.20.3) to map the MERFISH query dataset onto the extended single-cell lung cancer atlas (LuCA)^63^ reference dataset. In the reference, we select only cells belonging to normal, normal adjacent and primary tumor tissues. We merged the query and reference datasets, filtered out genes expressed in fewer than 10 cells and cells with fewer than 10 genes expressed, and performed CPM and log_2_ normalization. Initially, we ran scVI with default parameters, *technology* (MERFISH vs scRNA-seq) as a batch key, *assay* (MERFISH or the type of sc-RNA-seq assay) and *sample* as categorical covariates. The pre-trained scVI model was then used by scANVI to integrate the two datasets, leveraging the cell type annotations of the reference dataset, and to predict the cell type (including tumor cell) of the query cells. Manual curation was employed to correct errors in cell type assignment, predominantly in tumor cell detection.

For spatial clustering, we ran CellCharter with an L-neighborhood of 3 on the MERFISH dataset, using scVI with default parameters and *sample* as a batch key. Clustering stability analysis, using Fowlkes-Mallows Index and 10 repetitions between n = 2 and n = 25 clusters, revealed peaks at n = 5, n = 14, and n = 20. We opted for n = 20 for downstream analysis to highlight cellular niches containing various tumor subpopulations within the same sample. Cluster neighborhood enrichment, cell type enrichment, differential gene expression, and cancer cell signature score were analogously to the NSCLC CosMx dataset.

### Spatial clustering of Tissue Microarray lung cancer data

We ran CellCharter on a Tissue Microarray (TMA) dataset, composed of 416 lung cancer samples processed using Image Mass Cytometry (IMC) with 35 protein markers. Consistent with the original work, we excluded *GM-CSFR*, *PD-1*, *PD-L1* and *B7-H3* due to inconsistent staining and used raw IMC measurements. Given that no sign of batch effect between slides was present, no batch effect correction was performed. We set the number of clusters n = 30 as in the original work and the L-neighborhood to 3.

Focusing on the two spatial clusters with the highest enrichment of B cells (C15 and C17), using cell type labels provided by the authors, we fitted a Cox’s proportional hazard model to compute the association between the proportion of the two spatial clusters and the patient survival, with sex, BMI, smoking status, and stage as covariates. Three patients were excluded due to missing values.

We identified five spatial clusters as tumor-enriched (C0, C5, 13, 23, 29) because of their positive enrichment for tumor cells. Subsequently, we computed the asymmetrical cluster neighborhood enrichment between tumor-enriched and non-tumor-enriched clusters and observed that C23 was enriched in contacts with neutrophil-enriched clusters (C1, C7, C26). We identified markers for each of the tumor-enriched cluster by computing, for each marker, the mean z IMC intensity across tumor cells of that cluster, z-scored across all clusters.

### Association between hypoxia and neutrophil infiltration in bulk transcriptomic datasets

Gene sets of Tumor Associated Neutrophils (TAN) and Normal Adjacent tissue-associated Neutrophils (NAN) were sourced from a study characterizing immune cell infiltration in non-small cell lung cancer^63^. The hypoxia gene set was obtained from a study identifying 41 consensus cancer meta-programs across 24 tumor types (meta-program MP6)^97^. Bulk gene expression datasets of lung adenocarcinoma primary tumors were downloaded from public repositories reported in the respective studies^11,64–71^, along with clinical information. The TAN, NAN, and hypoxia signature scores for these bulk samples were computed using singscore^100^ v1.14.0 with default parameters. The association between these three continuous variables and overall survival was computed with Cox regression models implemented in the R package [survival](https://cran.r-project.org/web/packages/survival/index.html). Three models included each of the variables in isolation (without the other two); one model included all three variables. All four models also included and thus corrected for age, sex, and stage.

## DATA AVAILABILITY

The DLPFC Visium dataset was downloaded from the project’s Github page (https://github.com/LieberInstitute/HumanPilot). The mouse spleen CODEX dataset was obtained from the original publication^5^ (https://data.mendeley.com/datasets/zjnpwh8m5b/1). The NSCLC CosMx dataset was obtained from the original publication^41^ (https://nanostring.com/products/cosmx-spatial-molecular-imager/nsclc-ffpe-dataset/). The lung cancer MERFISH data was downloaded from Vizgen’s website (https://info.vizgen.com/ffpe-showcase). The lung cancer IMC dataset was obtained upon request to the corresponding author. The extended single-cell lung cancer atlas (LuCA)^63^ was downloaded from cellxgene (https://cellxgene.cziscience.com/collections/edb893ee-4066-4128-9aec-5eb2b03f8287). The mouse brain RNA+ATAC multi-omics dataset was downloaded from the UCSC Cell and Genome Browser (https://brain-spatial-omics.cells.ucsc.edu/). The lung cancer bulk RNA-seq datasets were retrieved from the original publications^11,64–71^ through the following links and Gene Expression Omnibus accession numbers: https://doi.org/10.5281/zenodo.3941896, https://gdac.broadinstitute.org/ (LUAD, rnaseqv2-RSEM_genes), GSE68465, [GSE72094](https://www.ncbi.nlm.nih.gov/geo/query/acc.cgi?acc=GSE68465), GSE31210, GSE50081, https://src.gisapps.org/OncoSG_public/ (GIS031), GSE37745, https://xenabrowser.net/datapages/ (Ding 2008).

## CODE AVAILABILITY

CellCharter is released as an open-source Python library on Github at https://github.com/CSOgroup/cellcharter. Code to reproduce all the analyses presented in this manuscript is available at https://github.com/CSOgroup/cellcharter_analyses.

## ACKNOWLEDGEMENTS

We thank R. Gottardo (Centre hospitalier universitaire vaudois, Lausanne), D. Gfeller (University of Lausanne), G. La Manno and E. Oricchio (École Polytechnique Fédérale de Lausanne) for their precious feedback during the development of this project. This project was in part supported by the Swiss Institute for Experimental Cancer Research TANDEM Research Project (G.C.). D.T. is supported by the Personalized Health and Related Technologies (PHRT) iPostdoc Project (grant no. 2022-476). A.S.M. is supported by the PHRT Technology transfer project (grant no. 2021-553).

## AUTHOR CONTRIBUTIONS

M.V. designed the algorithm and performed the computational analyses. D.T. analyzed the RNA-seq datasets. A.S.M. contributed to the interpretation of the CODEX mouse spleen results. L.A.W. provided the data for the IMC lung cancer dataset and contributed to the interpretation of the results. M.V. and G.C. designed the project and wrote the manuscript with contributions from all authors.

