## Supplementary Figures for "CellCharter reveals spatial cell niches associated with tissue remodeling and cell plasticity"

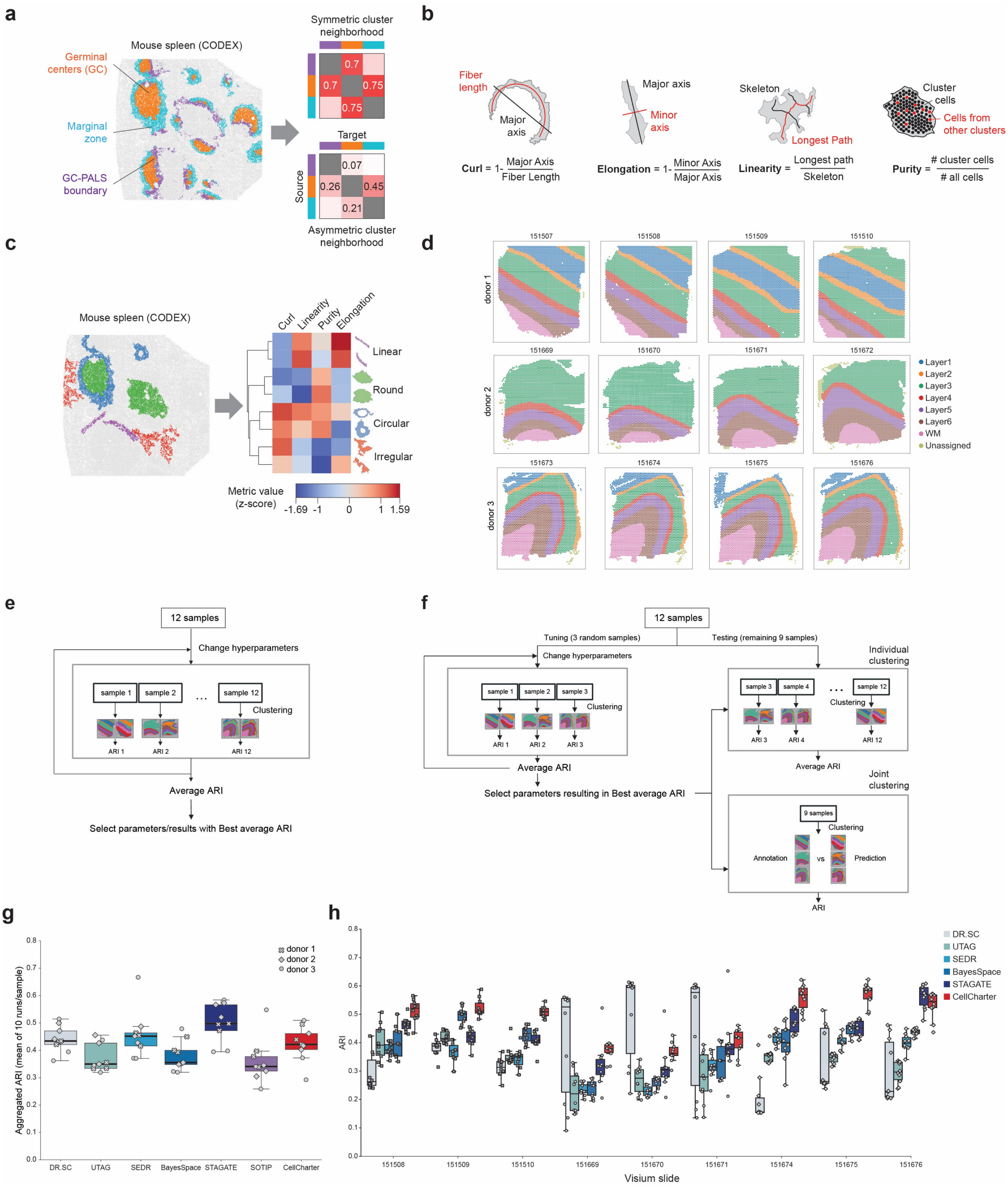


**Supplementary Figure 1: CellCharter's features and benchmarking of spatial clustering methods.**

**a)** Example of 3 spatial clusters (color coded) in a tissue sample of mouse spleen analyzed by CODEX (left) and symmetric (top right) vs. asymmetric (bottom right) neighborhood enrichment analysis. **b)** Schematic representation of the four metrics (curl, elongation, linearity, and purity) implemented in CellCharter to describe the shape of spatial clusters. **c)** Example of spatial cluster components (color coded) in a tissue sample of mouse spleen analyzed by CODEX (left). Heatmap representation of the shape metric values for each cluster component (right). Cluster components are grouped in representative shape classes: linear, round, circular, and irregular. **d)** Manual annotations of the Visium DLPFC samples. **e)** Previously adopted strategy based on hyperparameter tuning and clustering testing on the same dataset. **f)** strategy proposed in this study based on independent tuning and testing datasets. **g)** Mean Adjusted Rand Index (ARI) for each DLPFC sample (over 10 repetitions) obtained by the listed methods upon performing individual spatial clustering of the samples (n = 9 samples). The boxes show the quartiles of the dataset while the whiskers extend to points that lie within 1.5 inter-quartile ranges (IQRs) of the lower and upper quartiles. **h)** ARI for each DLPFC sample obtained by the listed methods upon performing joint clustering of all samples (n = 9 samples). The boxes show the quartiles of the dataset while the whiskers extend to points that lie within 1.5 IQRs of the lower and upper quartiles.

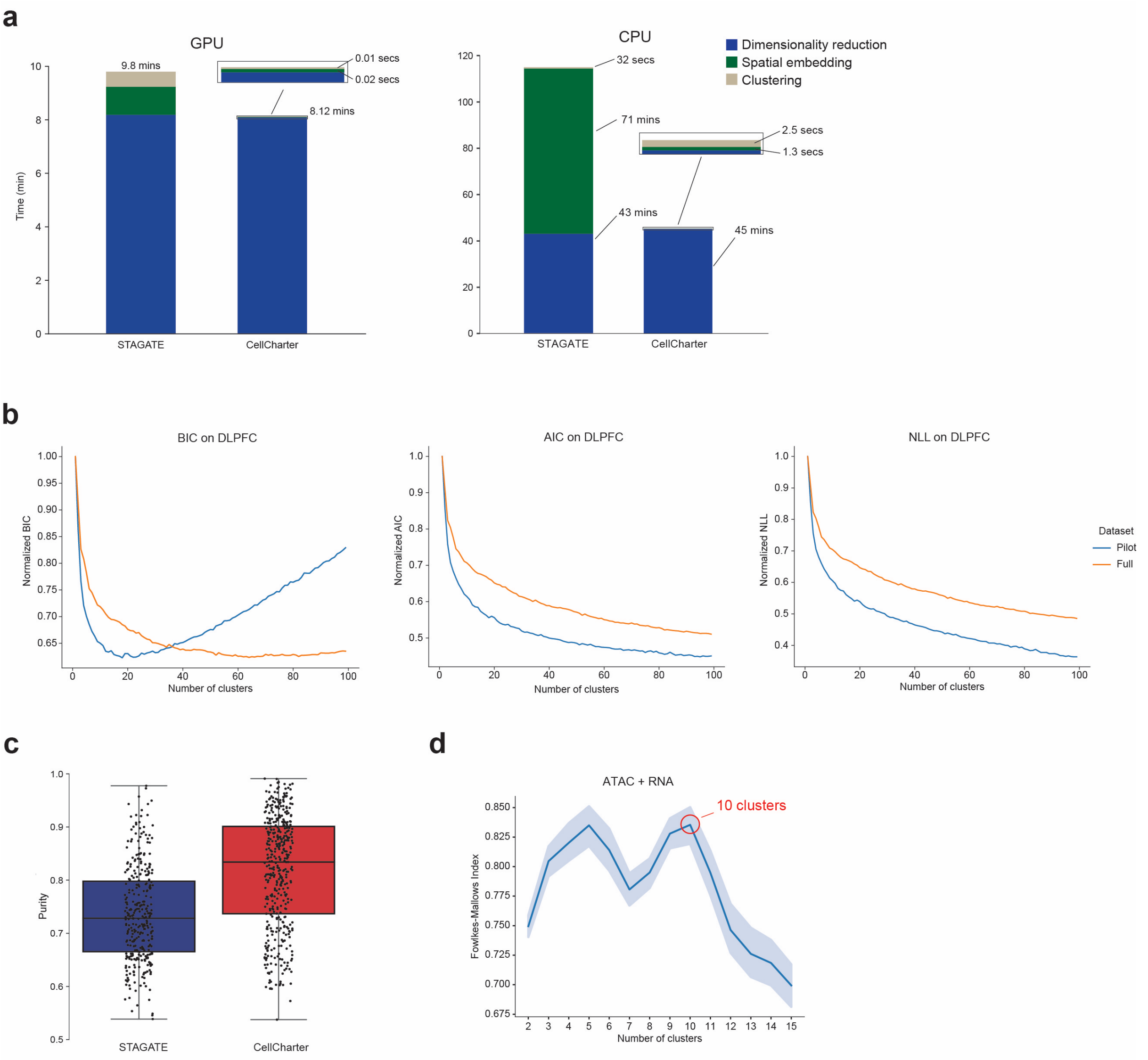


**Supplementary Figure 2: comparison between CellCharter STAGATE and evaluation of cluster stability analysis.**
**a)** Runtime divided by processing step of the two best-performing methods (STAGATE and CellCharter) in clustering all 12 samples of the DLPFC dataset on GPU and CPU. **b)** From left to right: Bayesian Information Criterion (BIC), Akaike Information Criterion (AIC), and Negative Log-Likelihood (NLL) values (y axis) for a number of clusters ranging between 2 and 100 (x axis) for the pilot (12 samples) and full (42 samples) version of the DLPFC dataset. **c)** Purity of the n = 756 cluster components in the CODEX mouse spleen dataset for the clusters of the two best-performing methods. The boxes show the quartiles of the dataset while the whiskers extend to points that lie within 1.5 inter-quartile ranges (IQRs) of the lower and upper quartiles. **d)** CellCharter cluster stability for range of numbers of cluster (x axis) using a concatenation of the embeddings from chromatin accessibility and gene expression. The most stable cluster solution is highlighted. Data presented as mean values (solid line) with a 95% confidence interval (shaded area).

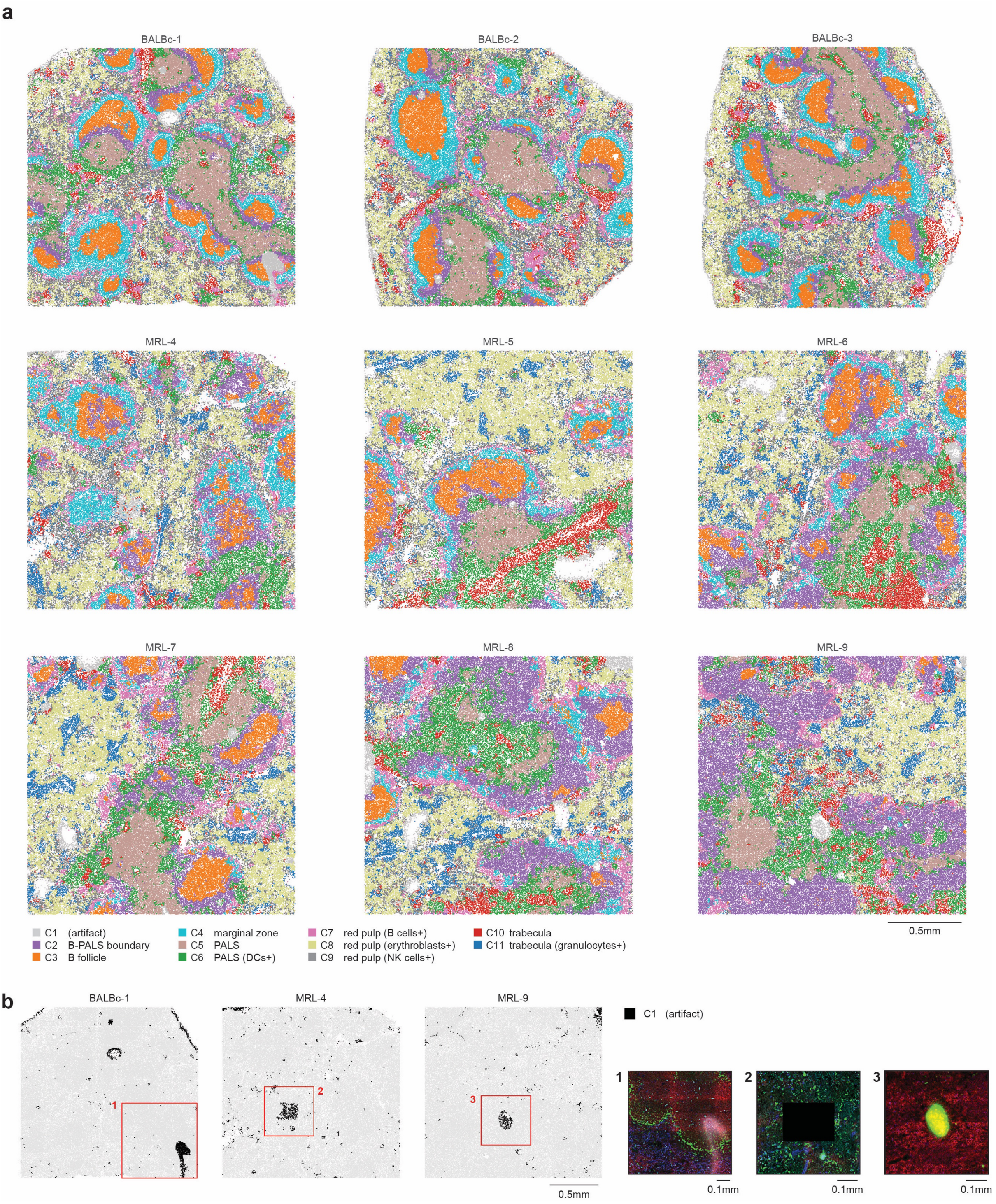


**Supplementary Figure 3: characterization of CellCharter's spatial clusters of the CODEX mouse spleen dataset.**

**a)** CellCharter's spatial cluster at n = 11 clusters for all cells of the 3 healthy (BALBc) and 6 systemic lupus erythematosus samples (MRL). **b)** (left) Spatial distribution of cells assigned to spatial cluster C1 in three selected samples. (right) Representative images of immunofluorescence staining artifacts corresponding to C1: autofluorescence (1), missing tile (2), fluorophore accumulation (3).

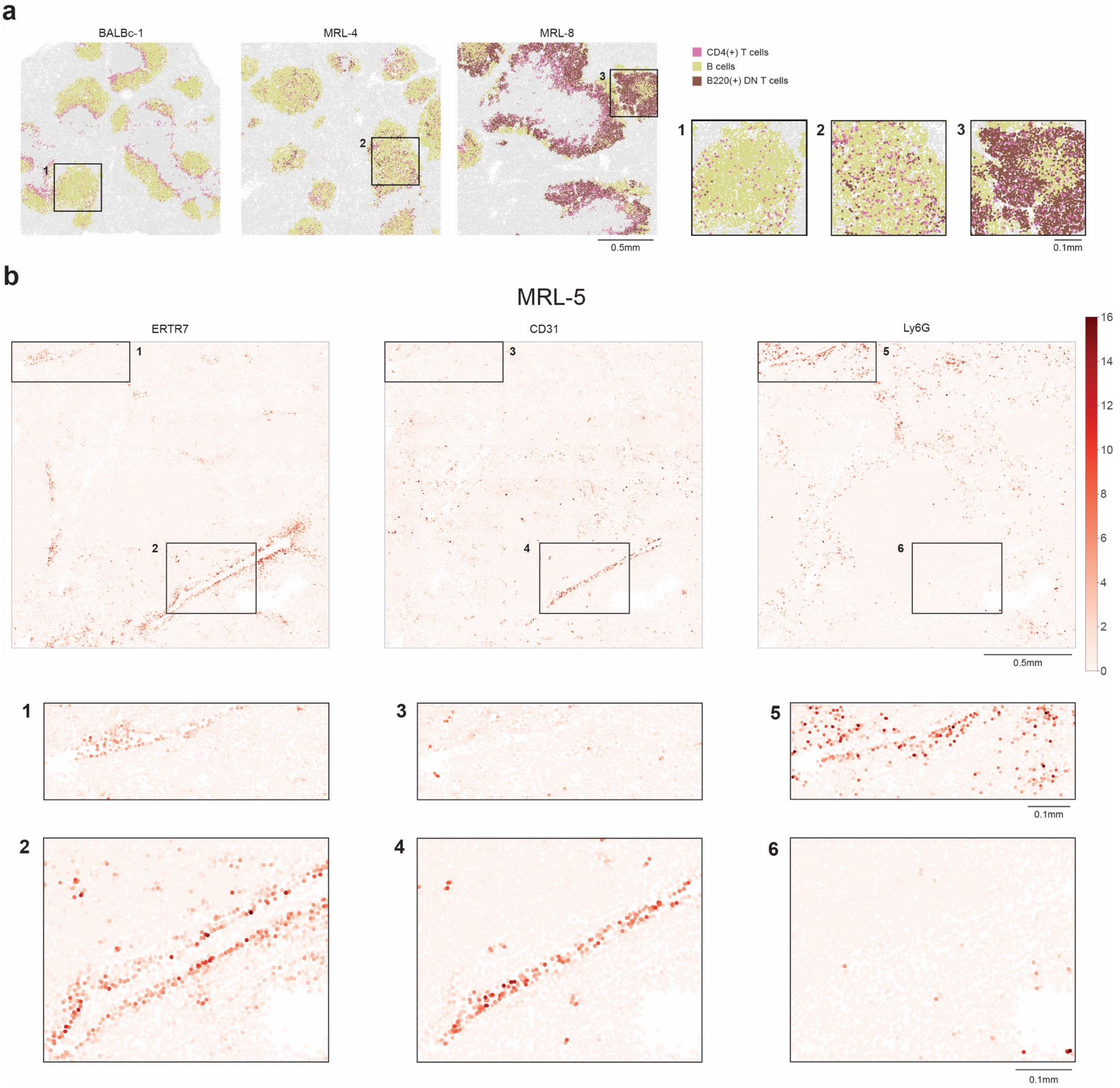


**Supplementary Figure 4: Differences in the spatial organization between healthy and systemic lupus spleen.**

**a)** Spatial distribution of $CD4^{+}$ T cells, B cells, and $B220^{+}$ T cells in the clusters associated to the germinal center, marginal zone, and GC-PALS boundary for representative examples of healthy (BALBc-1), early lupus (MRL-4) and intermediate lupus (MRL-8) spleen. **b)** Normalized intensity of *ERTR7*, *CD31*, and *Ly6G* markers in the intermediate lupus sample MRL-5. The MRL samples show the presence of two distinct clusters associated with trabecular structures. Insets (1 to 6) highlight the different expression of the markers in correspondence trabecular clusters C10 (insets 2, 4, 6) and C11 (insets 1, 3, 5).

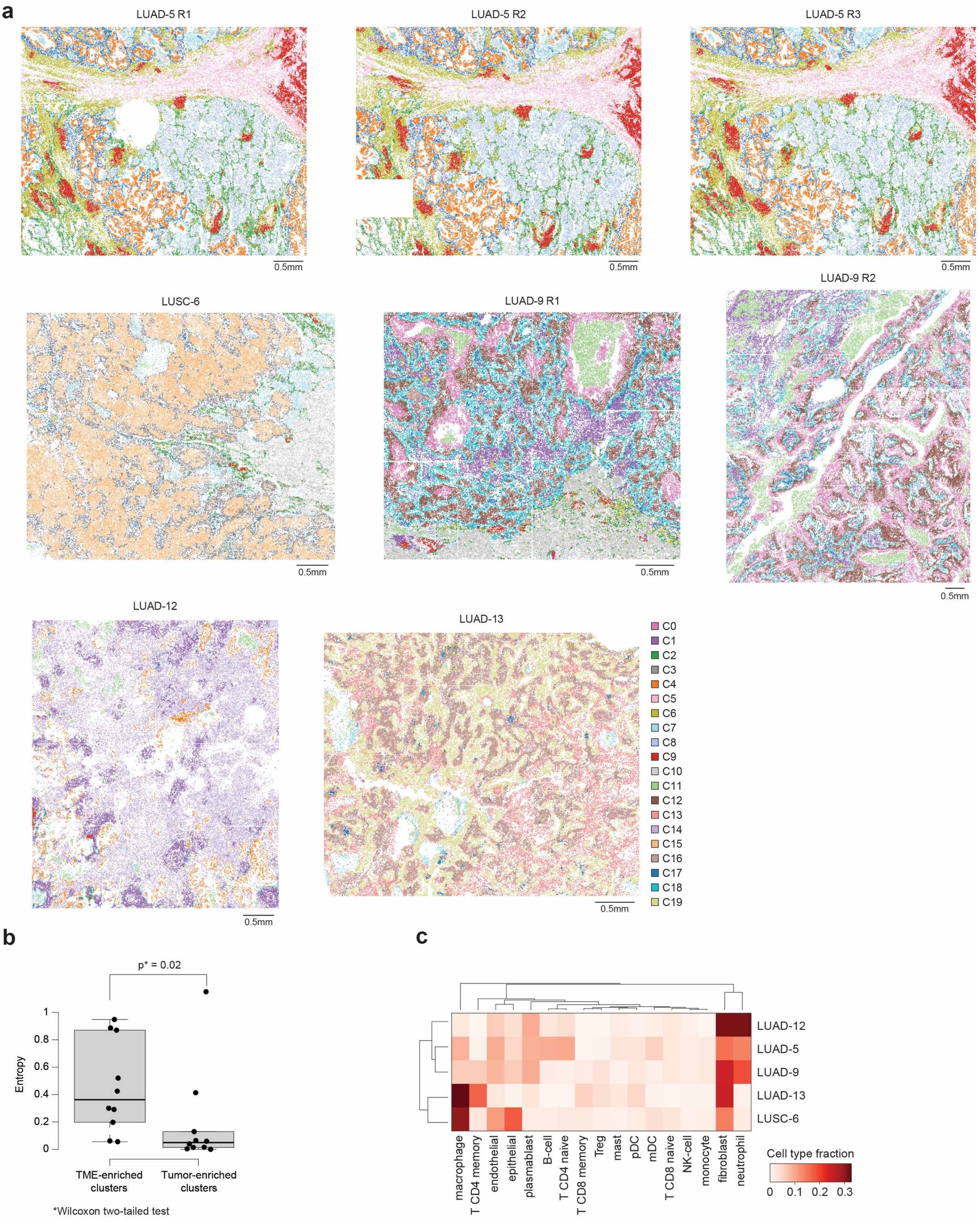


**Supplementary Figure 5: characterization of CellCharter's spatial clusters of the CosMx non-small cell lung cancer (NSCLC) dataset.**

**a)** CellCharter's spatial cluster at n = 20 clusters for all cells of the CosMx NSCLC samples. **b)** Entropy of patient proportion between n = 10 tumor microenvironment-enriched (TME-enriched) clusters and n = 10 tumor-enriched clusters. The thick central line of each box plot represents the median entropy, the bounding box corresponds to the 25th–75th percentiles, and the whiskers extend up to 1.5 times the interquartile range. p-values computed by two-tailed Wilcoxon test. **c)** Immune and stromal cell type fractions in tumor samples from 5 non-small cell lung cancer patients (LUAD: lung adenocarcinoma, LUSC: lung squamous cell carcinoma, NK-cell: natural killer cell).

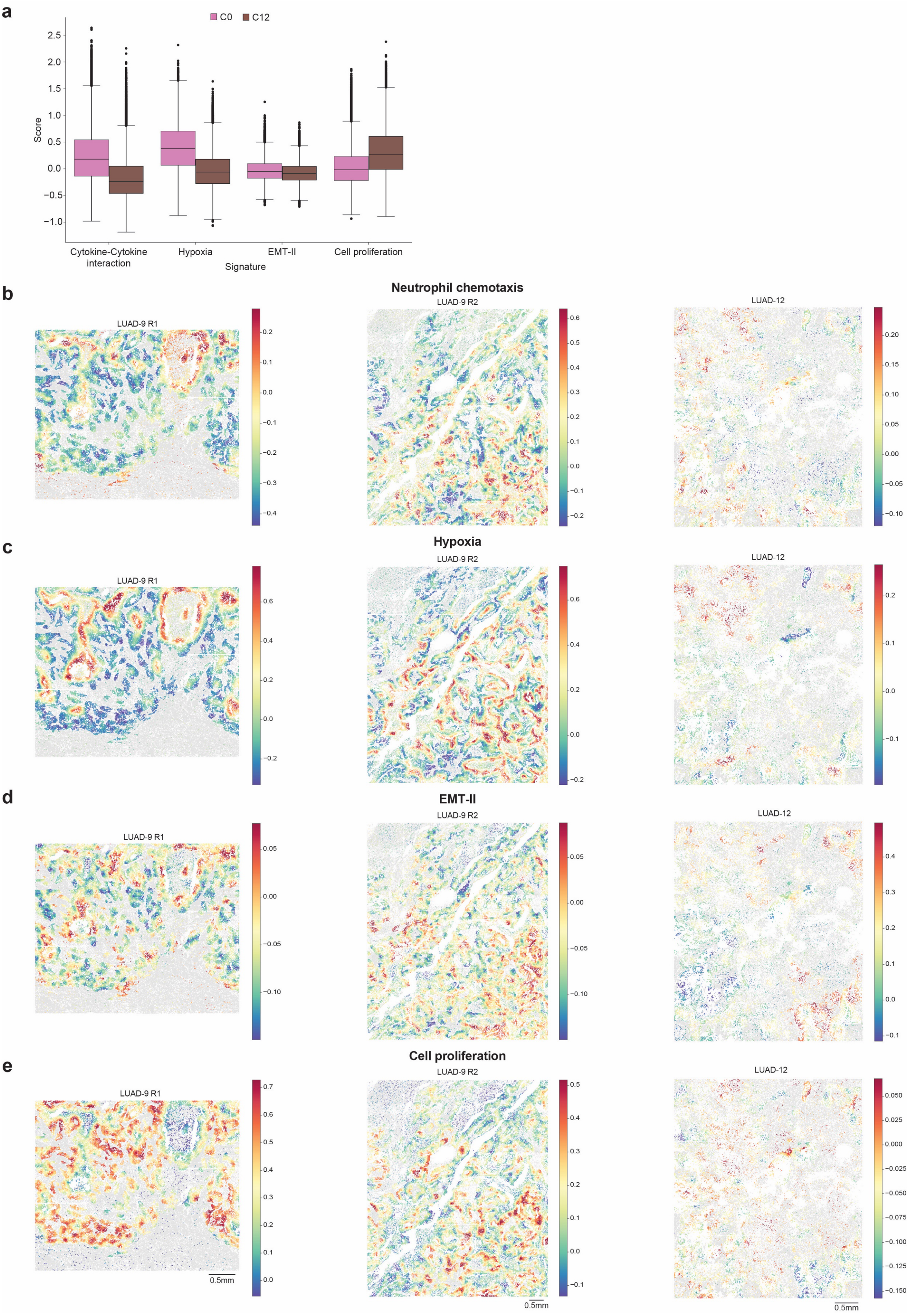


**Supplementary Figure 6: cell signature scores in two non-small cell lung cancer (NSCLC) patients.**

**a)** Gene expression signature score of n = 41,088 tumor cells in spatial clusters C0 and C12. The boxes show the quartiles of the dataset while the whiskers extend to points that lie within 1.5 inter-quartile ranges (IQRs) of the lower and upper quartile. **b-e)** Gene expression signature score of tumor cells of patients LUAD-9 and LUAD-12 for four gene signatures: neutrophil chemotaxis, hypoxia, epithelial-to-mesenchymal transition (EMT-II) and cell proliferation. The boxes show the quartiles of the dataset while the whiskers extend to points that lie within 1.5 IQRs of the lower and upper quartiles.

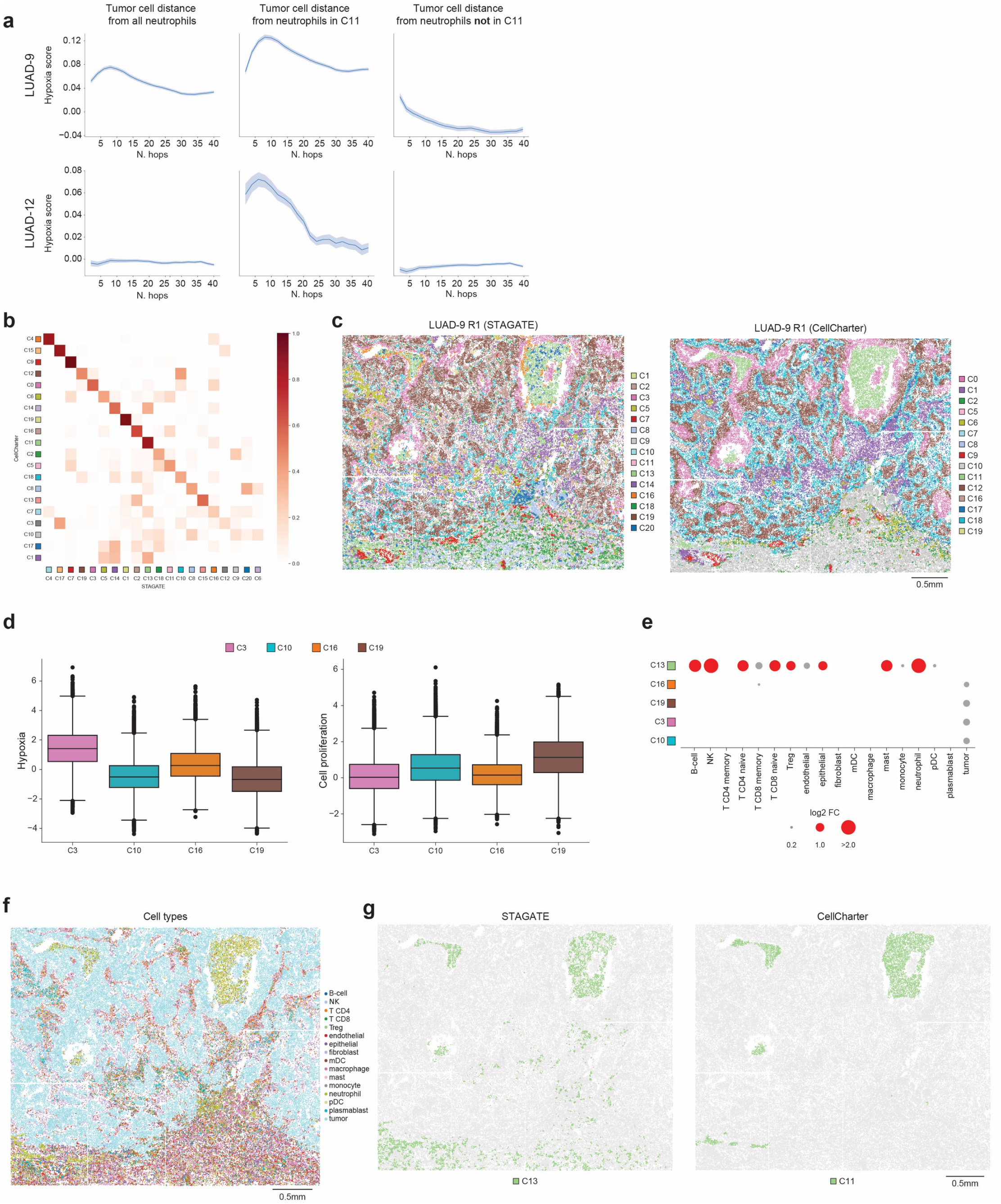


**Supplementary Figure 7: spatial organization of tumor associated neutrophils and hypoxic tumor cells in CellCharter and STAGATE.**

**a)** Average hypoxia gene signature score for tumor cells at increasing hop-distance from all neutrophils (left), neutrophils assigned to spatial cluster C11 (center), and neutrophils not assigned in cluster C11 (right). Error bands correspond to the 95% confidence interval. **b)** cluster concordance between CellCharter (rows) and STAGATE (columns). **c)** Example of CellCharter and STAGATE clusters on LUAD-9 R1. **d)** Hypoxia and cell proliferation signatures in n = 128,454 cells of tumor-enriched spatial clusters determined with STAGATE (NK: natural killer cell, Treg: regulatory T cell, mDC: myeloid dendritic cell, pDC: plasmacytoid dendritic cell). The boxes show the quartiles of the dataset while the whiskers extend to points that lie within 1.5 inter-quartile ranges (IQRs) of the lower and upper quartiles. **e)** Cell type enrichment of tumor-enriched clusters and neutrophil-enriched clusters obtained with STAGATE. **f)** Coarse cell type labels in LUAD-9 R1. **g)** Comparison of neutrophil-enriched clusters detected by STAGATE (left, C13) and CellCharter (right, C11).

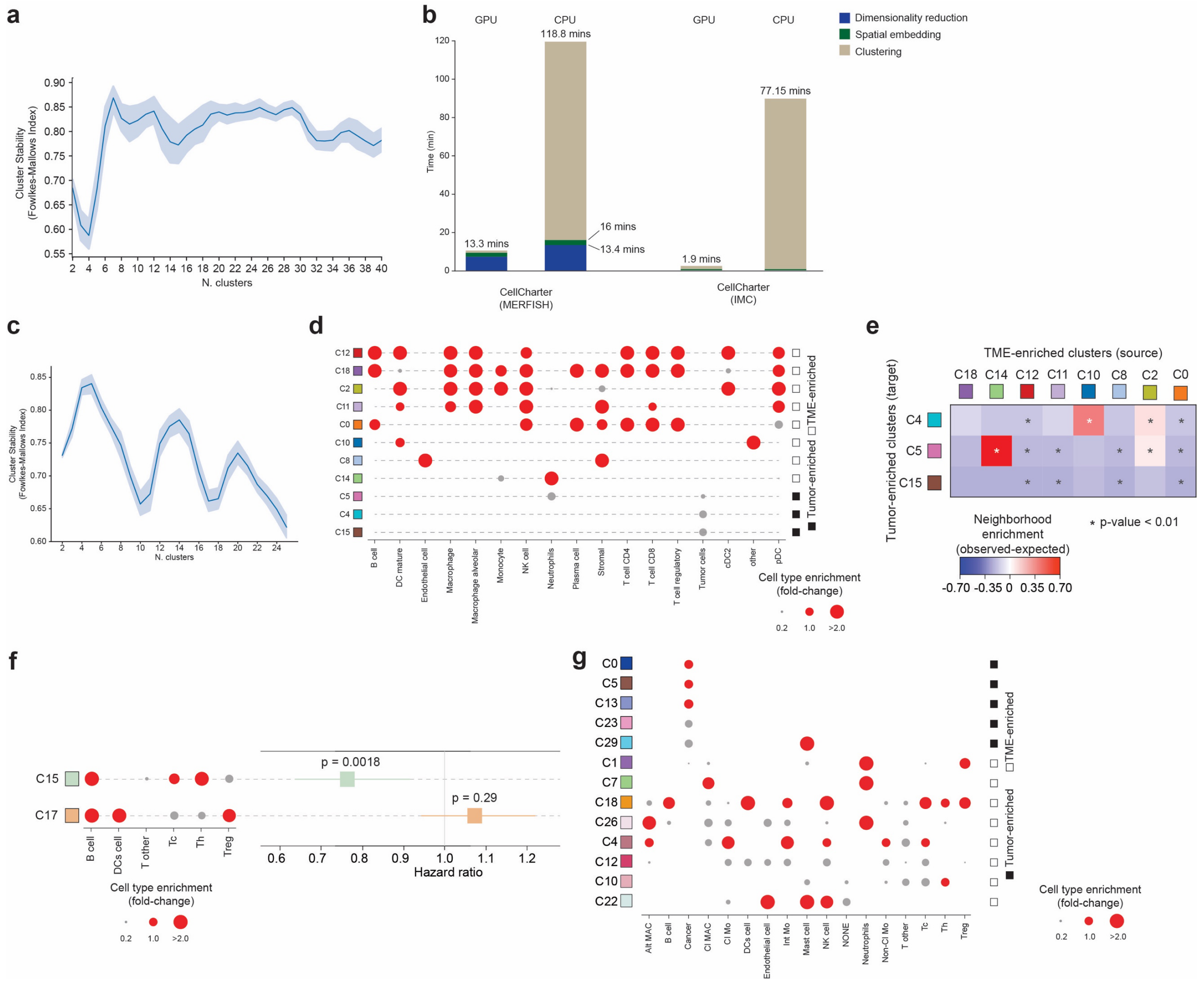


**Supplementary Figure 8: association between tumor hypoxia and neutrophil infiltration in the MERFISH and Image Mass Cytometry (IMC) datasets.**
**a)** Cluster stability (y axis) for range of numbers of cluster (x axis) on the full MERFISH dataset. Data presented as mean values (solid line) with a 95% confidence interval (shaded area). **b)** Runtime divided by processing step of CellCharter in clustering the MERFISH dataset (left) and IMC dataset (right) on GPU and CPU. **c)** Cluster stability (y axis) for range of numbers of cluster (x axis) on the 2 lung cancer samples of the MERFISH dataset. Data presented as mean values (solid line) with a 95% confidence interval (shaded area). **d)** Cell type enrichment of tumor-enriched and tumor microenvironment-enriched (TME-enriched) clusters in HumanLungCancerPatient1 obtained with CellCharter. **e)** Neighborhood enrichment between the TME-enriched clusters (source) and the tumor-enriched clusters (target) in HumanLungCancerPatient1. p-values computed by unpaired two-sided t-test. **f)** Cell type enrichment analysis (left) and hazard ratio and Cox-regression p-values (right) for the clusters C15 and C17 identified by CellCharter in n = 413 samples of the IMC dataset. Data are presented as mean values with 95% confidence interval. **g)** Cell type enrichment of tumor-enriched and TME-enriched clusters in the IMC dataset obtained with CellCharter. Only TME clusters for which there is a positive neighborhood enrichment with at least one tumor-enriched cluster are shown.

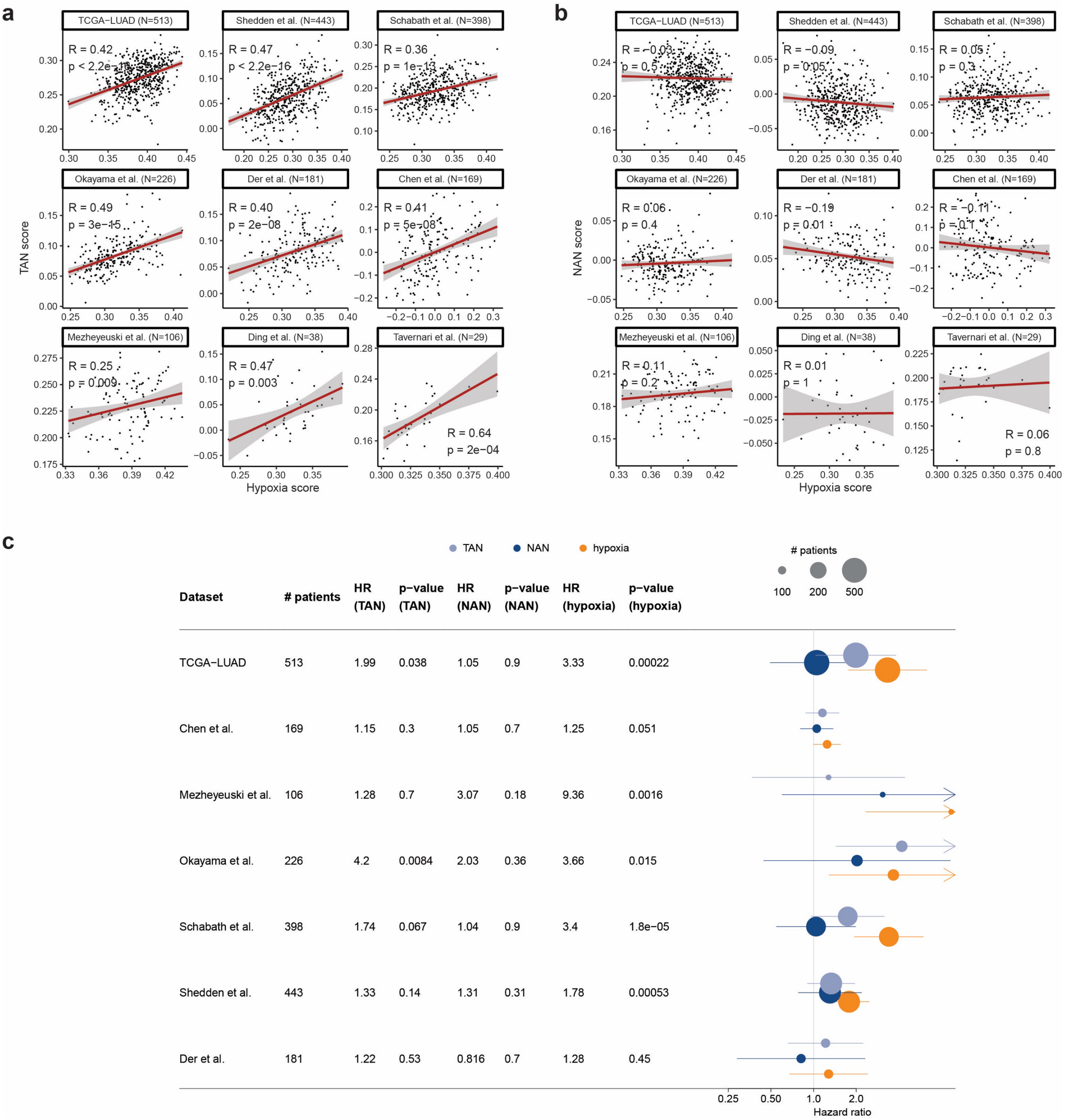


**Supplementary Figure 9: association of tumor-associated neutrophils (TANs), normal-associated neutrophils (NANs), and hypoxia with overall survival.**

**a)** Pearson correlation between hypoxia gene signature score and TAN score in 9 bulk RNA-seq datasets of lung adenocarcinoma. Data are presented as mean values (solid line) with a 95% confidence interval (shaded area). **b)** Pearson correlation between hypoxia gene signature score and NAN score in 9 bulk RNA-seq datasets of lung adenocarcinoma. Data are presented as mean values (solid line) with a 95% confidence interval (shaded area). **c)** Association of TAN, NAN, and hypoxia score with overall survival, corrected for sex, age, and stage, in the 7 bulk RNA-seq datasets that contained survival information. Data are presented as mean values with 95% confidence interval.
